# Spatial transcriptomic analysis of HIV and tuberculosis coinfection in a humanized mouse model reveals unique transcription patterns, immune responses and early morphological alterations

**DOI:** 10.1101/2025.01.29.635571

**Authors:** Sitaramaraju Adduri, Jose Alejandro Bohorquez, Rajesh Mani, Omoyeni Adejare, Diego Rincon, Joshua K. Kleam, Mounika Duggineni, Andy Omeje, Torry Tucker, Nagarjun V Konduru, Guohua Yi

## Abstract

*Mycobacterium tuberculosis* (*Mtb*) and human immunodeficiency virus (HIV) coinfection is one of the biggest public health concerns worldwide. Both pathogens are adept at modulating immune response and, in the case of *Mtb*, even inducing structural modification of the affected tissue. The present study aimed at understanding the early phenotypical and functional changes in immune cell infiltration in the affected organ, using a humanized mouse model. The humanized mice were infected with either HIV or *Mtb* in single infection, or with both pathogens in coinfection. Three weeks after the infection, lung samples were collected, and spatial transcriptomics analysis was performed. This analysis revealed high infiltration of CD4^+^ T cells in *Mtb* infection, but not in HIV or coinfection. Coinfected mice also showed a minimal number of NK cells compared to the other groups. In addition to infection status, histological features also influenced the immune cell infiltration pattern in the lungs. Two distinct airway regions with distinct immune cell abundance patterns were detected by spatial transcriptome profiling. A lymphoid cell aggregate detected in coinfection lung exhibited distinct transcript profile. The cellular architecture in the lymphoid cell aggregate did not follow the spatial patterns seen in mature granulomas. However, lymphoid cell aggregates exhibited granuloma gene expression signatures, and pathways associated with reactive oxygen species production, oxidative phosphorylation, and TGFβ and interferon signaling similar to granulomas. This study revealed specific transcription patterns, immune responses and morphological alteration signaling in the early stage of HIV and *Mtb* infections.

## Introduction

Tuberculosis (TB), caused by *Mycobacterium tuberculosis* (*Mtb*), continues to be a major public health concern. It has been estimated that as much as 25% of the world’s population is infected with *Mtb*, although most of these patients will develop the latent form of TB (LTBI)^1^. One of the main features of *Mtb* is its ability to modulate the immune response, facilitating the establishment of latent disease. This immune modulation occurs in circulating immune cells but also extends to resident tissue cells, where *Mtb* is able to induce structural modification in host tissue, forming specialized histological structures known as granulomas^2,3^. These structures are composed of a necrotic core enriched by diverse immune cell populations that contain the pathogen, preventing its spread throughout the host and the development of active TB. Consequently, there is growing interest in studying cellular infiltration in lung tissue during *Mtb* infection, particularly within granulomas^4–6^. Alterations in cellular subsets in the affected organ, coupled with intercellular communication, explain how these alterations either promote or hinder pathogen dissemination. Multiple microenvironments are present within granuloma structures that induce an immunoregulatory environment with differential signaling, characterized by pathway activation and cytokine production, which are critical for maintaining the granuloma structure^7^.

Local or systemic immune alterations can disrupt granuloma structures, facilitating bacterial growth^8,9^. The human immunodeficiency virus (HIV) is a major driver of the immune dysregulation, destabilizing granuloma structures that are critical for controlling *Mtb*^9^. In individuals coinfected with HIV in *Mtb*, the risk of developing active TB^10^ is significantly increased because of profound immunosuppression, primarily driven by the depletion of CD4*+* T cells, the main target for HIV infection, including those within granulomas. Reports show that CD4^+^ T cells within granulomas are particularly susceptible to HIV infection due to increased CCR5 coreceptor expression^8^. Furthermore, as CD4^+^ cells are essential for CD8^+^ cell maturation, the HIV-induced CD4^+^ cell depletion reduces the population of circulating effector and memory CD8^+^ cells, promotes immunosenescence in these cells, and impairs their functionality^11^.

Studies dedicated to understanding the cellular composition and intercellular networks present locally in the lungs during *Mtb* infection and HIV/*Mtb* coinfection have focused on infection models or samples in which infections and structural changes are already present in the host, leaving a gap in the knowledge of the early post-infection events^5–8,12^. This is partially due to the use of stored human samples or nonhuman primate (NHP) animal models for these experiments, which, while reliable, offer limited access to early infection timepoints. Recent studies have demonstrated the potential of humanized mice as effective models for studying *Mtb* and HIV^13,14^, capable of replicating immune changes caused by infection, including the lung granuloma formation. The present aimed to investigate early changes in the lung’s cellular repertoire induced by HIV, *Mtb*, or their coinfection, at an early timepoint in which granulomas have not been fully established. Additionally, we analyzed the transcriptomic changes in these cells using a spatial approach to determine alterations in cytokine production in the affected lung regions, enabling us to characterize the molecular processes that occur during infection and providing evidence of early granuloma development.

## Materials and methods

### Generation of humanized mice

Mice of the NOD.Cg-Prkdc^scid^ Il2rg^tm1Wjl^ Tg(CMV-IL3,CSF2,KITLG)1Eav Tg(IL15)1Sz/J (NSG-SGM3-IL15) strain (The Jackson Laboratory, Bar Harbor, ME), housed at the University of Texas Health Science Center at Tyler (UTHSCT) animal facility, were used for this study. Four-to five-week-old mice were irradiated at a dose of 100 cGy/mouse for myeloablation. Six hours later, each mouse was intravenously (IV) inoculated with 2×10^5^ human CD34^+^ hematopoietic stem cells (STEMCELL Technologies, Vancouver, Canada) for humanization.

At 16 weeks post-inoculation with human CD34^+^ cells, humanization was confirmed following an established protocol in our laboratory. Briefly, whole-blood samples were collected by puncture of the submandibular vein, and a density gradient was used to separate peripheral blood mononuclear cells (PBMCs). Once PBMCs were obtained, flow cytometry for specific human and mouse immune cell markers was performed, and humanization was confirmed in 13 mice. The criteria for humanization included a positive human/mouse leukocyte ratio, as determined by human CD45^+^ vs. mouse CD45^+^ expression, as well as the presence of multiple human immune cell subsets, as determined by markers for T cells (CD3^+^), B cells (CD20^+^), monocytes (CD14^+^) and natural killer (NK) cells (CD56^+^), as described in our published protocol^13^.

### Animal infection and experimental design

After confirming successful humanization, the animals were randomly assigned into 4 experimental groups (Figure 1A): A) no infection (n=3), B) HIV single infection (n=3), C) *Mtb* single infection (n=3) and D) HIV/*Mtb* coinfection (n=4). Animals in Groups C and D were infected with aerosolized *Mtb* or the H37Rv strain using a Madison chamber^13,15^. To verify the infection dose, three non-humanized mice were included in the infection chamber, euthanized the following day, and their lung samples were collected, macerated and plated on 7H10 plates supplemented with OADC (BD Biosciences, Franklin Lakes, NJ). These mice confirmed successful infection with an approximate dose of 40 CFU/lung.

**Figure 1.**
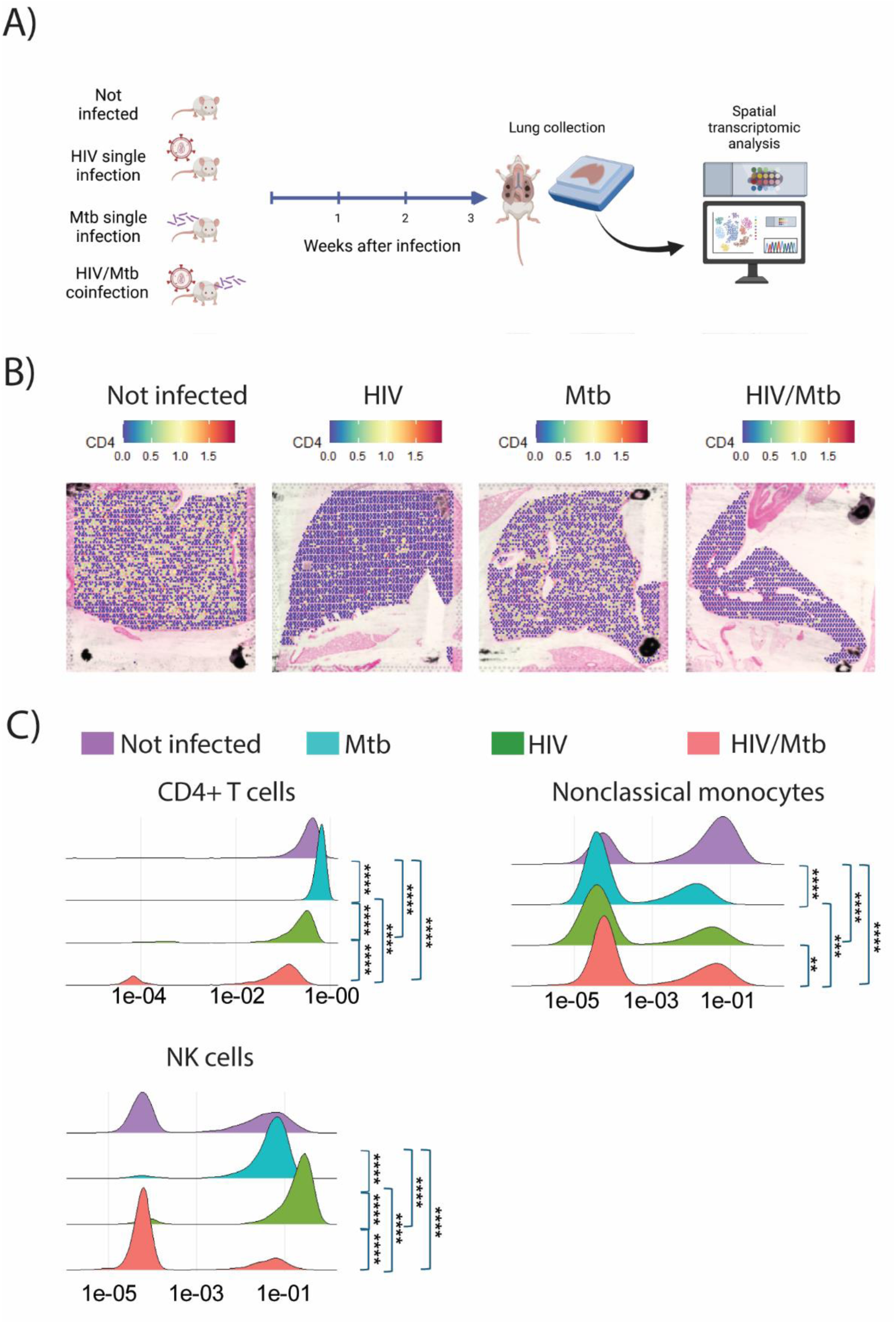
Cell type abundance variations between control, HIV, *Mtb* and coinfection mice as predicted by cell specific marker analysis and cell type deconvolution analysis. **A)** The experimental protocol. The number of mice was four per group. **B&C)** Expression levels of CD45 (PTPRC) and CD4 genes were shown across the four groups. **D)** Cell type weights for each spot in the samples based on RCTD analysis were plotted for each infection type. X axis shows cell types of weights which are proportional to the cell type abundance in the tissue. The difference in cell weight distributions was computed using Kruskal Wallis test followed by Dunn’s test for pairwise comparisons. P were adjusted for multiple hypothesis testing using Bonferroni correction method. Significance levels are indicated as \**P* < .05, \*\**P* < .01, \*\*\**P* < .001, and \*\*\*\**P* < .0001.

Two days after *Mtb* infection, the mice in Groups B and D were intraperitoneally (IP) injected with 10^7^ TCID_50_ of the HIV-1 BaL strain (obtained from the NIH AIDS Reagent Program). Two weeks after inoculation, whole blood samples were obtained, and RNA was extracted from the plasma samples using a NucleoSpin RNA isolation kit (Macherey-Nagel, Allentown, PA) as per the manufacturer’s instructions. Viral RNA load in each animal was quantified using RT–qPCR with a control standard of known HIV-1 genome copy numbers (obtained from the NIH AIDS Reagent Program), as previously described^16^. All HIV-inoculated mice were positive for HIV via RT‒qPCR and presented similar viral RNA loads.

Three weeks after infection, all the animals were humanely euthanized in accordance with the NIH guidelines for the euthanasia of rodents with carbon dioxide, with confirmation by cervical dislocation. During necropsy, lung samples from each mouse were collected in accordance with previous protocols. Briefly, once euthanasia was confirmed, the thoracic cavity was opened, and the lungs were perfused and inflated by injection of 10% neutral buffered formalin (Thermo Scientific, Waltham, MA) into the right heart ventricle and trachea, respectively. Afterward, the samples were immediately submerged in the same buffer and kept at 4 °C until paraffin embedding^17,18^. All animal procedures were approved by the UTHSCT Institutional Animal Care and Use Committee (IACUC) (Protocol #763).

### 10x Visium spatial transcriptome assay

Formalin-fixed, paraffin-embedded (FFPE) tissue sections were used to prepare sample slides according to the 10x Genomics protocol (CG000518 and CG000520). RNA was extracted from the FFPE sections using the RNeasy FFPE Kit (Qiagen, 73504) according to the manufacturer’s protocol. Samples with DV200 values > 30% were selected for spatial transcriptomics assay. After hematoxylin and eosin (H&E) staining and imaging, qualified samples were decrosslinked per protocol (CG000520), and the sequencing libraries were prepared according to the manufacturer’s protocol (CG000495). The quality of the final libraries was assessed using the Agilent High Sensitive DNA Kit (5067-4626) with an Agilent Bioanalyzer 2100 (Agilent Technologies, Santa Clara, USA). The qualified libraries were sequenced using paired-end sequencing on an Illumina NovaSeq System (Illumina, Inc., USA).

### Visium spatial sequencing data analysis

With demultiplexed sequence fastq files and CytAssist tissue images as inputs, space ranger was used to map the reads to the latest human genome (hg38) and quantify the read counts per spot. The counts were normalized via the ‘sctranform’ method in the Seurat package^19^. For cross-sample comparisons, data were further normalized using the minimum median count function. The samples were clustered and projected on UMAP. The “Find marker” function was used to identify cluster-specific markers and differentially expressed genes, defined as genes with a minimum |log2FC| value of 2 at an adjusted p value of <0.05. Pathway analysis was performed using GprofileR^20^. The Loupe browser was used to examine the spot-wise expression levels of individual genes. Deconvolution of spatial gene expression profiles was performed using the RCTD package^21^. Single-cell RNA sequencing data (GSE192483) of lung tissues with 18F-FDG avidity and nearby uninvolved tissues from six tuberculosis patients^22^ were used as a reference for deconvolution analysis. Azimuth^23^ was used for annotating the cells in the reference dataset. Visualizations were created using the ggplot2 package in the R environment (4.1.1). Gene set enrichment analysis (GSEA) was performed using the ‘fgsea’ package with default settings, leveraging C8 datasets from molecular signatures database. The gene sets for human were downloaded into R environment using ‘msigdb’ package. Differentially expressed genes identified by ‘FindMarkers’ function of ‘seurat’ were ranked based on average logFC and adjusted p values were used as input for GSEA. GSEA was performed using ‘fgsea’ command with 1000 permutations on the ranked list of genes. An adjusted p value of 0.25 was cut off as suggested by the original documentation of the GSEA.

### Imaging mass cytometry (IMC)

IMC was performed on 5-micron thick sections from FFPE blocks. The sections were baked at 60^0^C for 2 hours and dewaxed in xylene for 20 minutes. The sections were rehydrated in a graded series of ethyl alcohol followed by water. The sections were incubated at 96^0^C for 30 minutes in 1x target retrieval solution with pH 9 (Cat #S236784-2, Agilent) and cooled at 70^0^C for 20 minutes. The sections were blocked with 3% bovine serum albumin in PBS for 45 minutes at room temperature. An antibody cocktail was applied, and the slides were incubated overnight at 4℃ in a hydration chamber. After washing with 0.2% Triton X-100 and with PBS, the sections were counterstained with Ir-Intercalator (cat# 201192A; Standard Biotools) at 1:400 dilution in PBS for 30 min at room temperature. The sections were air dried at room temperature for 20 minutes. Imaging was performed on Hyperion XTi imaging system (Standard Biotools). Regions of interest were selected using CYTOF software, and the data was analyzed using MCD viewer.

### Immunofluorescent staining

Formalin-fixed paraffin-embedded (FFPE) lung sections from *Mtb* single-infected and *Mtb*/HIV-coinfected mice were processed for immunofluorescence staining as follows. Tissue sections-containing slides were incubated at 60 °C for 2 hours to melt the paraffin, followed by complete deparaffinization in fresh xylene (4 × 5 min each). Sections were then rehydrated through a graded ethanol series (100%, 95%, 70% and 50%; 10 min each) and rinsed in distilled water. Antigen retrieval was performed by incubating slides in 10 mM sodium citrate buffer (pH 6.07) at 90 °C for 1 hour. After cooling to room temperature, nonspecific binding was blocked using the protein concentrate and mouse Ig blocking reagent from the M.O.M. Immunodetection Kit (Vector Laboratories). Sections were then incubated with primary antibodies against surface markers for human CD68 and CD3 (Invitrogen, Thermofisher) and intracellular markers for IRF-7 (Invitrogen, Thermofisher) and TGF-β (Santa Cruz) overnight at 4 °C. The following day, slides were washed and then incubated with anti-mouse secondary antibodies conjugated with AF488, AF568, and AF647 (Invitrogen, Thermofisher) for 2 hours at room temperature. After washing, coverslips were mounted with DAPI-containing medium, and images were acquired on a Zeiss LSM-500 confocal microscope. All micrographs were processed using ImageJ (Fiji, NIH).

## Results

### Immune cell infiltration patterns in the lungs vary distinctly across each infection type

We generated humanized mice using the NSG-SGM3-IL15 mouse strain. As expected, the expression of human IL-3, GM-CSF, and KITLG genes led to the development a full-lineage of human immune cells, similar to our previously reported humanized NSG-SGM3 mice^24^. Additionally, the transgenic expression of the human IL-15 gene resulted in a substantial portion of differentiated human natural killer (NK) cells (**Figure S1**). Using this humanized NSG-SGM3-IL15 mouse model, we investigated the effect of HIV and *Mtb* coinfection on immune responses in the lungs. In our prior study on a humanized mouse model of coinfection, we observed rapid CD4^+^ T-cell depletion within 15 days post-infection and plateaued after 28 days. To examine immune responses during the rapid decline of CD4^+^ cells, we euthanized mice 21 days post-infection and performed spatial transcriptome analysis of lung specimens from humanized mice with HIV infection, *Mtb* infection, and coinfection, using the 10x Genomics Visium platform. The uninfected group served as a control (**Figure 1A**). After excluding the poor-quality regions and empty spots, our spatial transcriptomics analysis detected the expression of 18,000 genes across 3633, 2698, 2778, and 1397 spots in uninfected, HIV-infected, *Mtb*-infected, and coinfected lungs, respectively.

As TB primarily affects lungs, analyzing the infiltration frequency of various immune cells into lungs is critical to understand the immune response against *Mtb*. CD4^+^ T cells play crucial roles in controlling *Mtb* infection by releasing signals that activate B cells to produce antibodies, cytotoxic T cells to eliminate infected cells, and macrophages to engulf and destroy pathogens. Consequently, we sought to investigate how reduced CD4^+^ T cell levels impact immune cell infiltration in lung tissues during *Mtb*/HIV coinfection. To compare the abundance of immune cell markers across all 4 infection groups, we normalized spatial gene expression data from all samples using the sctranform method and then corrected the minimum median counts of the SCT data to ensure comparability across samples. As shown in **Figure 1B**, expression of CD45 (PTPRC gene) is elevated in lung tissue from *Mtb* single infection compared to all other infection groups. CD45 is expressed in all nucleated lymphoid cells and over expression of CD45 indicates excessive immune cell infiltration in *Mtb* lungs compared to others. Next, we observed minimal CD4 expression in the HIV single infection and coinfection groups compared with the uninfected and *Mtb* single infection groups, consistent with HIV-driven massive depletion of CD4^+^ T cells (**Figure 1C**). These results suggest an infection-specific landscape of immune cell infiltration in the lungs.

To further investigate immune cell infiltration frequencies in the lungs across different infection groups and controls, we performed RCTD-based cell type deconvolution analysis by integrating previously published single-cell RNA sequencing data of human lung granulomas (as detected by PET-CT) and their matched normal tissues from TB patients with our spatial transcriptome data from all 4 infection groups: uninfected, HIV-only, *Mtb*-only, and *Mtb*/HIV-coinfected. We first annotated cell types in the scRNA-seq data using the Azimuth annotation database at the finest annotation level and then transferred these labels to our spatial transcriptome data during deconvolution analysis in ‘full mode’. Given that Visium spots may span up to 6 individual cells, we opted to run deconvolution in ‘full mode’, as it allows identification of any number of cells existing in each pixel. Consistent with CD4 gene expression in the tissues, cell type weights for the CD4^+^ T cells also revealed the highest abundance of CD4^+^ T cells in the *Mtb*-infected mice, with reduced abundance in the HIV-infected and coinfected mice (**Figure 1D**). In line with prior reports, CD4^+^ T cell abundance was lower in coinfected mice compared to those with HIV-only infection^25^ (**Figure 1D**). Unexpectedly, the abundance of nonclassical monocytes was lower in all infection groups compared to uninfected control (**Figure 1D**). NK cell abundance was lower in the coinfection group than in the other three groups, and the NK cell abundance was the highest in the lungs with *Mtb* single infection (**Figure 1D**), corroborating earlier findings of reduced NK cell infiltration during coinfection relative to *Mtb*-only infection^25^. The distributions of the weights of myeloid lineage cells, such as alveolar macrophages, CCL3^+^ alveolar macrophages, classical monocytes, monocyte-derived macrophages, and interstitial macrophages, were similar across all groups, suggesting that infection status has minimal or no effect on the infiltration of these cells (**Figure S2A**). However, B cells exhibited reduced abundance in HIV-infected and coinfected groups compared to uninfected and *Mtb*-only infected controls (**Figure S2B**). Taken together, these results suggest that lung tissue infiltration of various immune cell types depends on the type of infection.

### Validation of lung immune cell infiltration patterns using IMC

To validate the results of spatial transcriptome data, we performed IMC on lung tissues from one mouse with *Mtb*-only infection and one with *Mtb*/HIV coinfection. IMC is a multiplexed imaging technique that enables comprehensive and accurate analysis at the single-cell level with spatial resolution using metal tagged antibodies. IMC confirmed a reduced abundance of CD4+ T cells in coinfected lungs compared to *Mtb*-only infection (**Figure 2A**), supporting the findings of spatial transcriptome data. Similarly, the frequency of CD8+ T cells (**Figure 2B**) and B cells (**Figure S3A**) were less in coinfection group compared to *Mtb*-only infection. While robust deconvolution of spatial data was limited due to the underrepresentation of certain cell types in the reference dataset, IMC enabled analysis of cell population not covered in the spatial data deconvolution. Notably, granulocytes, which were not assessed in spatial transcriptomics analysis due to their absence in the reference dataset used for deconvolution, showed a lower abundance in coinfected lungs compared to *Mtb*-only infection (**Figure 2C**). Similarly, dendritic cells also showed lower abundance in coinfection compared to *Mtb*-only infection (**Figure S3B**). Collectively, both spatial transcriptomics and IMC consistently demonstrated reduced infiltration of major immune cell types in confected lungs compared to *Mtb*-only infection, suggesting a weakened immune response against *Mtb* in the context of HIV infection.

**Figure 2:**
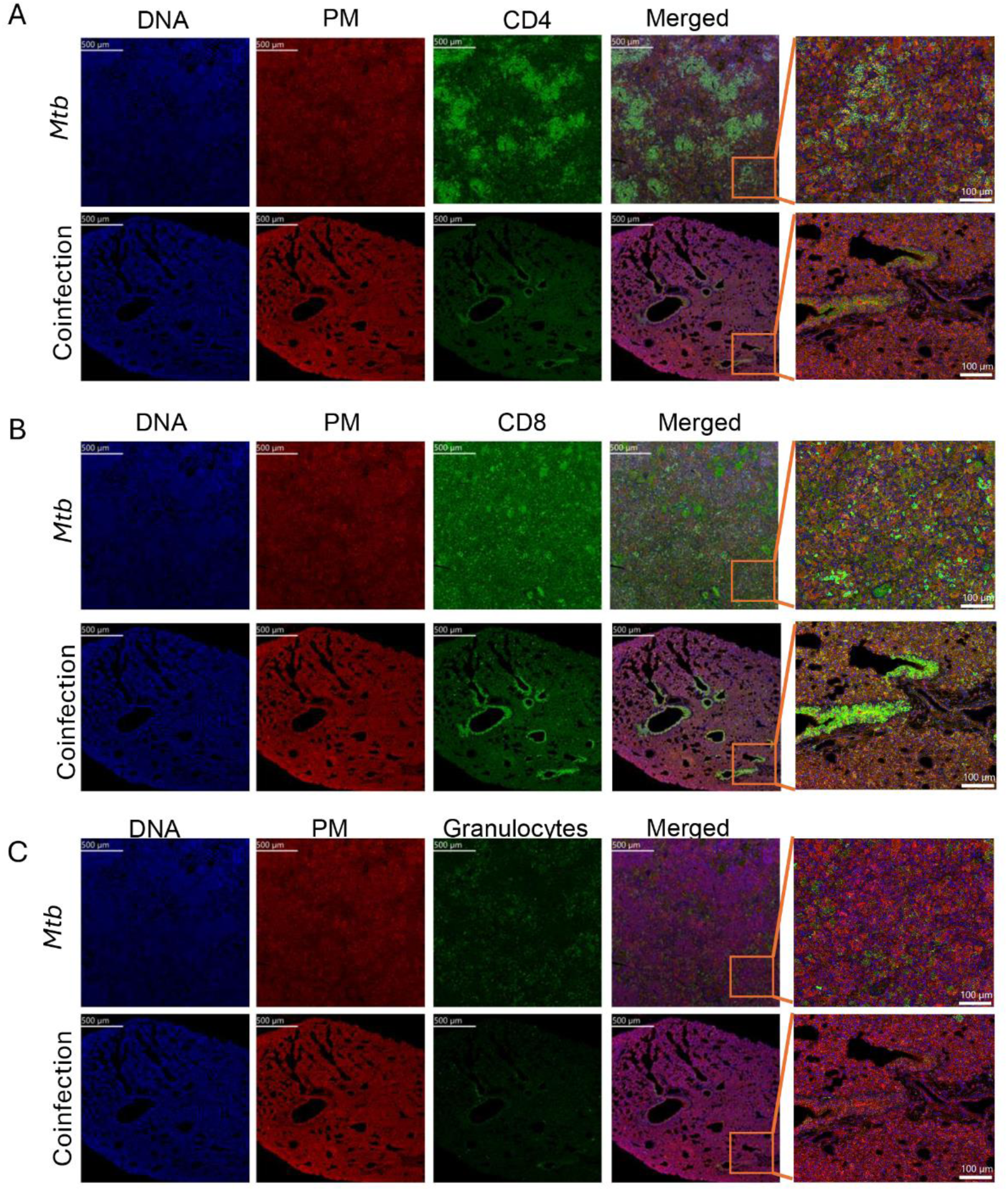
**Spatial distribution map of CD4+ and CD8+ T cells, and granulocytes in the lungs from *Mtb* infection and coinfection mice**. Ir-Intercalator was used to stain nucleus. 195Pt_ICSK2 was used for staining plasma membrane. 156Gd_CD4, 162Dy_CD8, and 160Gd_CD66b were used to detect CD4+ T cells **(A)**, CD8+ T cells **(B)**, and granulocytes **(C)**, respectively.

### Lung immune cell infiltration patterns differ depending on infection status and the local histological milieu

To understand the distinct patterns of immune cell infiltration into the lungs, we clustered the Visium spots across all 4 infection states together using uniform manifold approximation and projection (UMAP) based on 50 principal components at a resolution of 0.9. This analysis identified a total of 14 clusters across 4 samples: no infection (12), HIV (11), *Mtb* (12), or coinfection (12) (**Figure 3A, S4A**). Clusters 0-11 were present in all 4 samples, whereas Cluster 12 was absent in HIV and coinfection samples. Intriguingly, Cluster 13 was detected exclusively in the coinfection group (**Figure S4A**).

**Figure 3.**
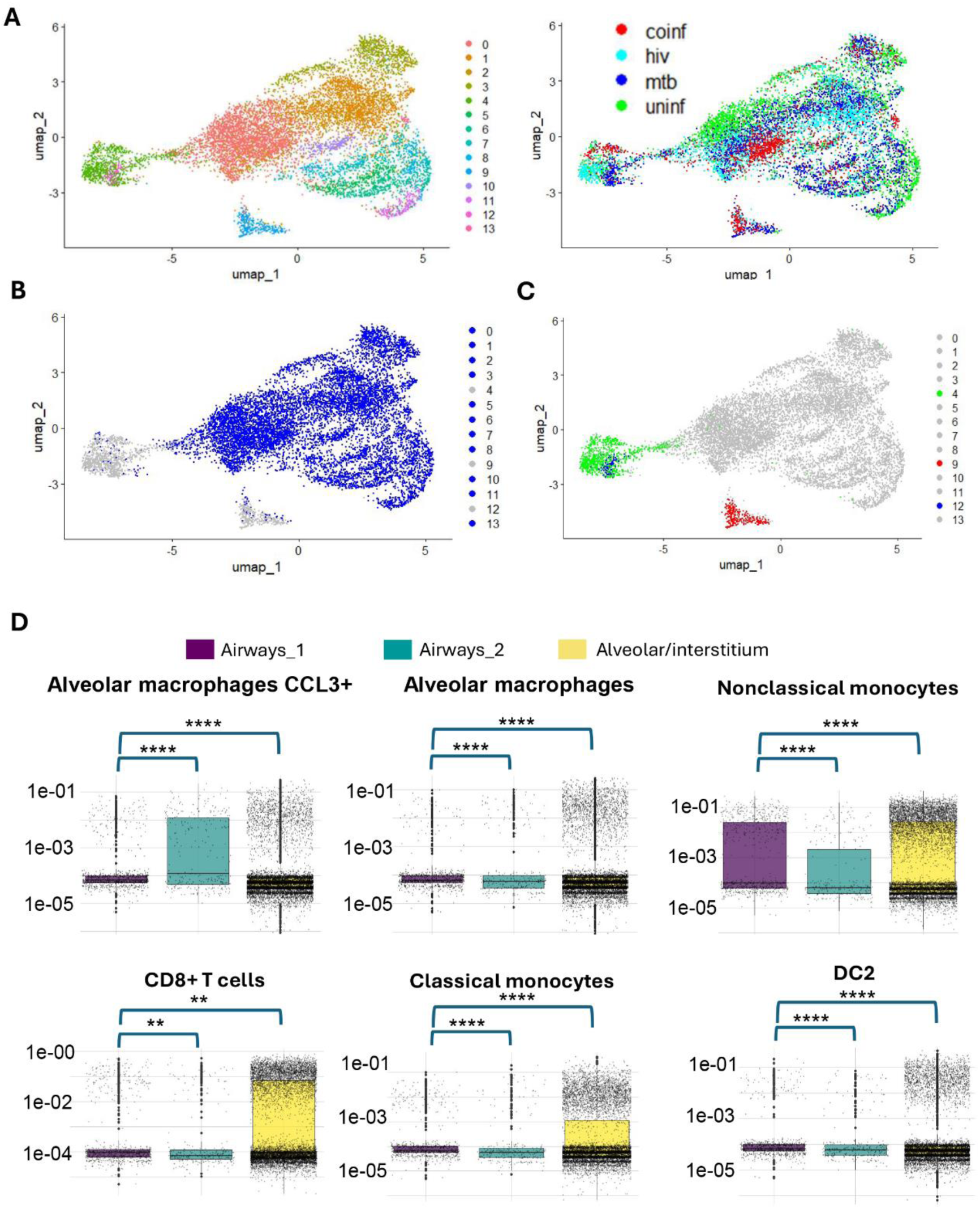
Identification clusters based on gene expression of all tissue spots from no infection, HIV, *Mtb*, and coinfection mice lungs using Seurat package. **A)** Clusters from all four samples projected on UMAP and color based on cluster identifier (left) and sample type (right). **B)** Alveolar/interstitium cluster (shown in blue), **C)** airway_1 (shown in blue/green) and airway-2 (shown in red) clusters on UMAP projection. **D)** Cell type weights obtained in RCTD deconvolution analysis were plotted for each cluster. Y axis shows cell type weights which are proportional to the cell type abundance in the corresponding cluster. The color key indicated the cluster identifier. The difference in cell weight distributions was computed using Kruskal Wallis test followed by Dunn’s test for pairwise comparisons. P values were adjusted for multiple hypothesis testing using Bonferroni correction method. Significance levels are indicated as \**P* < .05, \*\**P* < .01, \*\*\**P* < .001, and \*\*\*\**P* < .0001.

The UMAP analysis (**Figure 3A**) revealed a broad separation of all 14 clusters into three distinct groups (**Figure 3A**). The majority of the clusters, including Clusters 0-3, 5-8, 10, 11 and 13 (11 total clusters), were placed together on the right of the UMAP projection as a single large group (**Figure 3B**). These spots corresponded to the alveolar/interstitial compartment of the lung across all 4 infection categories (**Figure 3B, S4B**). Therefore, we combined these 11 clusters into one large cluster, namely, the ‘alveolar/interstitial’ cluster, for further analysis. Clusters 4 and 12, positioned separately and distant from the alveolar/interstitial cluster on the UMAP projection (**Figure 3C**; represented in green and blue, respectively), were closer to each other than to cluster 9 on UMAP projection, suggesting similarities between cluster 4 and 12. (**Figure S4C**; represented in green and blue, respectively). Therefore, we combined Clusters 4 and 12 into one cluster called ‘airway_1’ (**Figure S4C**; represented in red) for preliminary analysis. Similarly, Cluster 9, another distinct cluster far away from the rest on the UMAP, also harbored spots representing airways across all four groups. It was then renamed ‘airway_2’ for further analysis. Despite both airway-1 and airway_2 clusters representing airways, they were not combined due to their distinct separation on the UMAP. Interestingly, cluster separation was driven by tissue location and histological features rather than infection status. These findings suggest that immune infiltration is more strongly influenced by histological features than by infection status in this study.

To investigate the differences in immune infiltration patterns across histological clusters, we analyzed cell type weights obtained from the deconvolution analysis described in Figure 1. Monocyte-derived macrophages were more abundant in the airway-1 cluster compared to other groups (**Figure S4D**). Notably, the airway-2 cluster was enriched with CCL3^+^ alveolar macrophages, which are known for effective bacteria and antigen phagocytosis (**Figure 3D**)^26^. In contrast, alveolar macrophages with lower CCL3 expression were slightly more abundant in the airway_1 cluster (**Figure 3D**). Additionally, nonclassical monocytes were less abundant in the airway_2 cluster and alveolar/interstitium clusters compared to the airway_1 cluster (**Figure 3D**).

The alveolar/interstitial cluster was enriched with CD8^+^ T cells and classical monocytes, whereas the number of type 2 dendritic cells was lower (**Figure 3D**). The abundance of B cells, mast cells and type 1 dendritic cells was similar across the three clusters (**Figure S4E**). These results suggest that different histological regions may harbor distinct immune cell types. However, it remains unclear whether this differential abundance contributes to distinct immune responses.

### Airways of *Mtb* infection exhibit different immune cell patterns compared to HIV and coinfection

In uninfected mice, all airways exclusively belonged to cluster 4. In contrast, mice with HIV and coinfection have both cluster 4 and cluster 9, but not cluster 12. The cluster 12 was unique to *Mtb*-only infected mice. Given that cluster 4 is present in uninfected mice, and clusters 9 and 12 appear to be stratified based on infection type, we compared the transcriptomic signatures in cluster 12 and 9. As our spatial assay was specific to human transcriptome, the robust segregation of two different airway clusters and their stratification by infection type suggests that the airways in *Mtb*-only infected group may harbor a unique immune cell population compared to those with HIV or coinfection. To investigate this further, we performed Gene Set Enrichment Analysis (GSEA) using human immunologic gene sets (C8, n=886) to map the immune response signatures associated with both clusters. We sought to decipher how HIV infection affects airway immune cell population and differs from *Mtb* infection. GSEA analysis on clusters 9 and 12 revealed a reduced abundance of proinflammatory M1 macrophages in HIV infection (**Figure 4A**), which produce TNF-alpha and IL-12 and exhibit anti-mycobacterial activity through nitric oxide production and autophagy. Highlighting the disruption of immune response against *Mtb* in coinfection, GSEA revealed the diminishing of M1 macrophages in cluster 9 which represents airways from HIV and coinfection. This depletion of M1 macrophages would potentially facilitate the progression of *Mtb* infection. GSEA also revealed lower abundance of neutrophils (**Figure 4B**) and classical monocytes (**Figure 4C**), alongside increased frequency of NK cells (**Figure 4D**), innate lymphoid cells type 2 (**Figure 4E**), and nonclassical monocytes (**Figure 4F**). Taken together, these results indicate that immune cell populations at the airway interfaces may vary across disease type and HIV infection may disrupt the immune cell infiltration patterns that aid the survival of TB pathogen.

**Figure 4.**
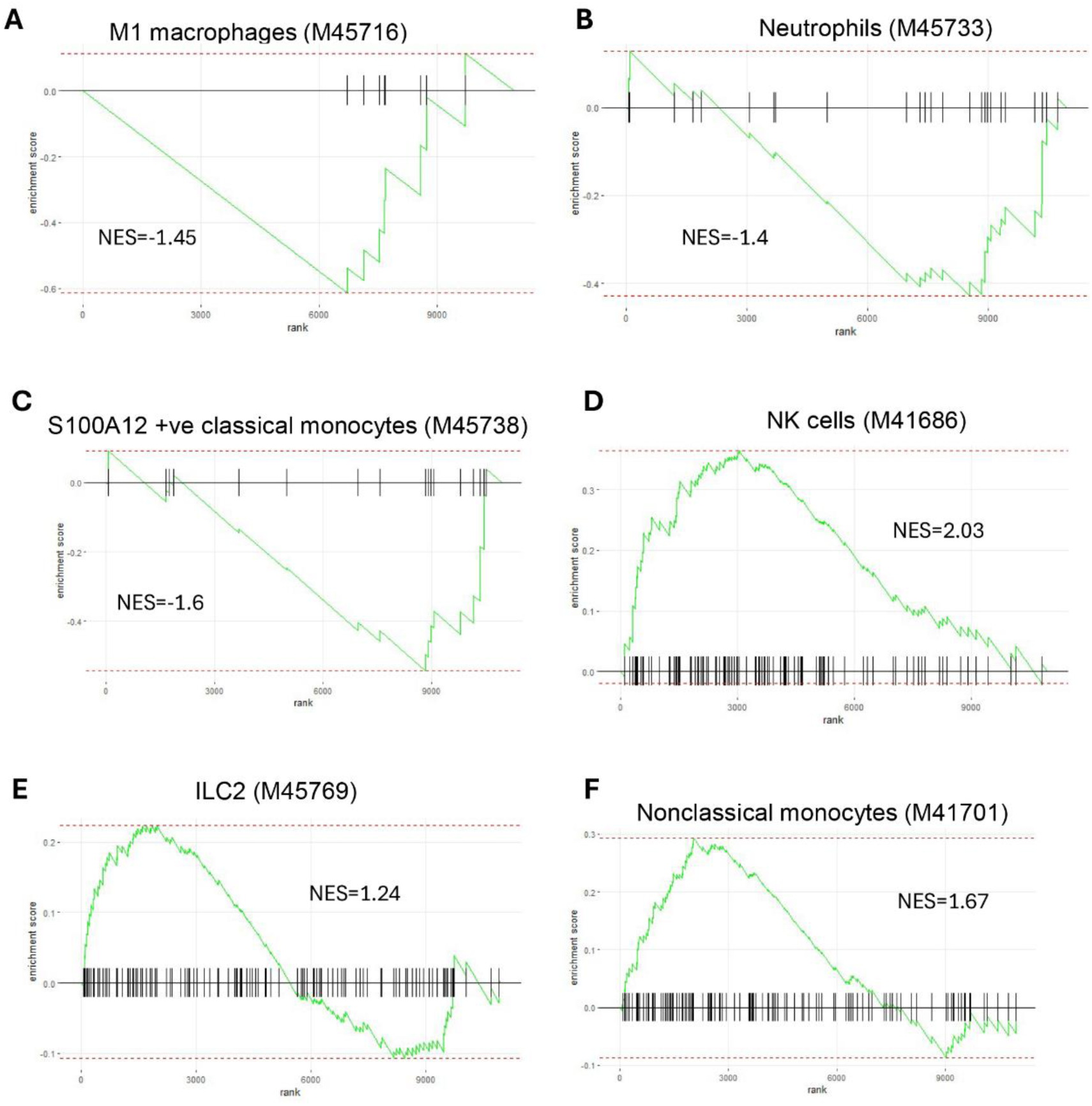
Gene set enrichment analysis for identifying differences in immune cell types between cluster 9 compared to cluster 4 shown in. figure 2C. The enrichment plots illustrate the enrichment of the corresponding gene set in the ranked list of genes based on comparison between Cluster 9 vs cluster 4. Gene set names for each enrichment plot were shown on the top. **A)** M1 macrophages, **B)** Neutrophils, **C)** S100A12 +ve classical monocytes, **D)** NK cells, **E)** type 2 Innate lymphoid cells, and **F)** nonclassical monocytes. The x-axis represents the rank of genes based on their differential expression, ordered from most upregulated to most downregulated. The y-axis shows the running enrichment score (ES), which quantifies the degree to which the gene set is overrepresented at the top or bottom of the ranked list. The green curve depicts the cumulative ES, with peaks indicating maximum enrichment. Vertical black lines mark the positions of genes within the gene set. The Normalized Enrichment Score (NES) reflects the enrichment score normalized for gene set size and dataset variability, with positive/negative values indicating enrichment in upregulated/downregulated genes in cluster 9, respectively. Msigdb identifier of the gene set corresponding to each enrichment plot was shown in parenthesis.

### Cellular organization and composition of lymphoid cell aggregates markedly differ from those of typical mature tuberculosis granulomas

Several previous studies have investigated the cellular architecture of mature tuberculosis granulomas^12,27–29^. A granuloma may appear as a tiny lymphoid cell aggregate at the initial stages and progress into a nonnecrotic or central necrotic mature granuloma. However, our understanding of the cellular architecture of lymphoid cell aggregates remains limited compared to that of mature granulomas. Hematoxylin (H&E) staining identified a lymphoid aggregate in the region of tissue from coinfected mice subjected to spatial transcriptome profiling. Focusing exclusively on this region, we investigated two key questions: 1) whether the cellular architecture of lymphoid aggregates resembles or differs from that of mature granulomas, and 2) whether the lymphoid aggregates present any transcriptomic changes characteristic of mature granulomas.

Remarkably, we observed that spots belonging to Cluster 13, the only UMAP cluster that was exclusively detected in coinfection, spanned the entire lymphoid cell aggregate in this sample, indicating a unique transcription profile distinct from other tissues across all 4 infection groups (**Figure 5A**). This lymphoid aggregate, located in the interstitial region adjacent to a large airway, appears to have densely infiltrated lymphoid cells on H&E staining (**Figure 5B**). On UMAP, Cluster 13 co-localizes with other interstitial region clusters but remains distinctly separate from them (**Figure 5C**).

**Figure 5:**
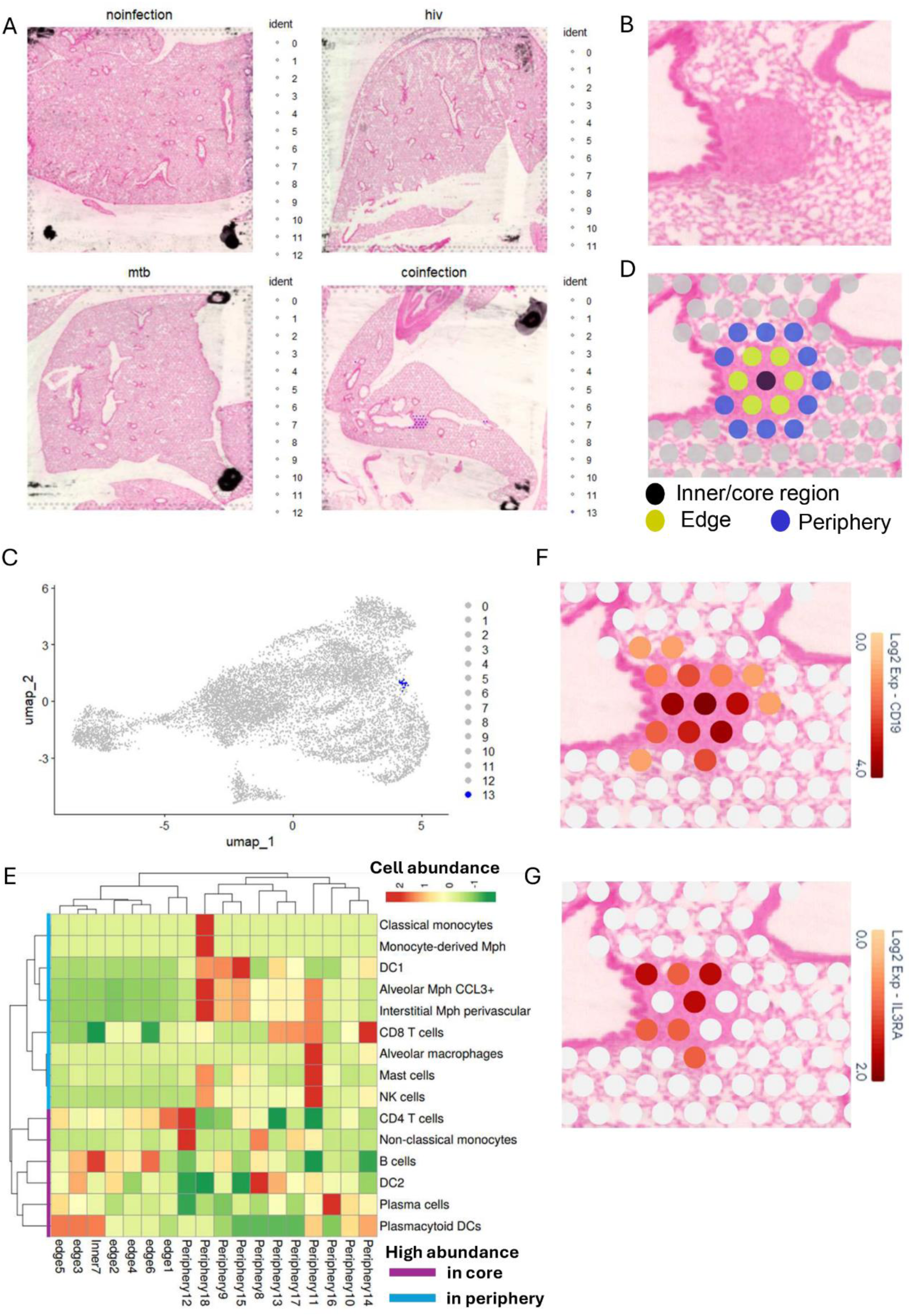
Characterization of the unique cluster (cluster 13) detected in HIV/*Mtb* coinfection mouse lung. **A)** Spatial mapping of cluster 13 which was detected exclusively in coinfection lung. Clusters were identified on UMPA projection of all Visium spots passing quality control from all four infection groups combined. Spots representing cluster 13 were highlighted in blue. **B)** Hematoxylin staining of lung region matching with cluster 13 location in coinfection lung. This region contains a circular lymphoid aggregate formed by dense lymphocyte infiltration located within the interstitial region and close to two airways. **C)** UMAP projection of all Visium spots passing quality control from all four infection groups highlighting the location of cluster 13. **D)** Projection of the Visium spots overlapping with lymphoid aggregate shown in Figure 3B. The spots were labeled as core/inner region, edge, and periphery depending on the location and were highlighted according to the label with different colors. **E)** Heatmap showing the cell type abundances all the spots highlighted in Figure 3D. Columns show individual spots spanning the lymphoid aggregate. The rows show the immune cells detected in the spots. Color intensities represent the abundance of each cell in each spot. Row and column dendrograms represent the similarity between cell abundances and individual spots, respectively, as computed using hierarchical clustering. **F)** Spatial map of expression of CD19, a mature B cell marker within and around the lymphoid aggregate. Spot-wise gene expression data was projected on hematoxylin staining image of the lung tissue. Color key represents the level of transcript expression. **G)** Expression of CD123 (IL3RA), plasmacytoid markers in the same region as described for Figure 5F.

Necrotic granulomas are structured around the central core of necrotic cell debris in which many bacteria are concentrated. This necrotic core is surrounded by layers primarily composed of macrophages^2^, including epithelioid macrophages, interspersed with classical alveolar macrophages, monocyte-derived macrophages, lipid-laden foamy macrophages, and giant cells^2^. A variety of other immune cell types are observed at the periphery and within the epithelioid layers of the granuloma. Non-necrotic granulomas have similar structural arrangements except that they do not have necrotic cores^2^. However, the cellular architecture and dynamics of these lymphoid aggregates are poorly understood.

To decipher the cell type abundances in the Visium spots spanning the lymphoid cell aggregate, we extracted cell type frequency predictions obtained from RCTD deconvolution analysis. We labeled spots spanning the lymphoid cell aggregate as inner/core regions, edges and peripheries, beginning from the center to the outer region (**Figure 5D**). To compare the abundance of different immune cells predicted using deconvolution analysis, we plotted a heatmap with hierarchical clustering of genes and spots (**Figure 5E**). Interestingly, we observed that the spotwise clustering, as depicted in the column dendrogram, effectively distinguished the peripheral spots from the inner and edge spots, indicating location-based stratification of cell distribution. Unlike mature tuberculosis granulomas, where B cells predominantly present in the periphery and are absent or scarce in the core, lymphoid aggregates exhibited a high abundance of B cells in the inner/core region (**Figure 5E**). This was supported by elevated expression of mature B-cell markers, such as CD19 (**Figure 5F**), CD20 (MS4A1) (**Figure S5A**), and IgM (IGHM) (**Figure S5B**). Plasmacytoid dendritic cells (pDCs) were also highly enriched in the core and edge regions, as shown in the heatmap (**Figure 5E**), consistent with high expression of pDC markers such as CD123 (IL3RA) (**Figure 5G**) and CD303 (CLEC4C) (**Figure S5C**). Similarly, lymphoid aggregates abundantly harbor nonclassical monocytes (**Figure 5E**), marked by elevated levels of CD97 (ADGRE5), P2RX1, and Siglec10^30^. As expected, we observed a high abundance of CD97 (ADGRE5) (**Figure S5D**), P2RX1 (**Figure S5E**), and Siglec10 (**Figure S5F**) in the core or edge of the lymphoid aggregate, with reduced abundance in the periphery. Additionally, we also observed enrichment of plasma cells, DC2s, CD4+ T cells and nonclassical monocytes in the core of the lesion (**Figure 5E**). In contrast, NK cells, mast cells, macrophages, CD8+ T cells, DC1 cells, and classical monocytes were predominantly found in the periphery (**Figure 5E**).

Taken together, these results suggest that lymphoid cell aggregates may not essentially originate with macrophages at their core. We hypothesize that granuloma architecture evolves over time, developing into a macrophage-laden core surrounded by a lymphocyte-abundant periphery.

### Lymphoid cell aggregates displayed gene expression profiles typical of granulomas

Given that lymphoid cell aggregates do not show a cellular architecture typical of a mature granuloma, we assessed whether this lesion exhibited any transcriptomic changes associated with mature granulomas. We examined the expression of genes that are part of gene sets related to granulomas using the Molecular Signatures Database (MSigDB). We analyzed the HP_GRANULOMA gene set, which includes ACP5, CYBB, CYBC1, DNASE2, DOCK2, IRF8, NF1, and PIK3CG. The lymphoid aggregate region showed higher aggregated expression of these genes compared to other lung tissues (**Figure 6A**), with individual genes also exhibiting elevated expression in this region (**Figure S6A**). Similarly, another gene set, HP_ GRANULOMATOSIS, consisting of ASAH1, CTLA4, CYBA, CYBB, HLA-DPA1, HLA-DPB1, NCF1, NCF2, PRTN3, and PTPN22, also exhibited increased expression in lymphoid aggregates compared to the surrounding tissue (**Figure 6A, S6A, S7A**). These results suggest that gene expression changes characteristic of mature granulomas emerge early during granuloma development from the lymphoid aggregate.

**Figure 6:**
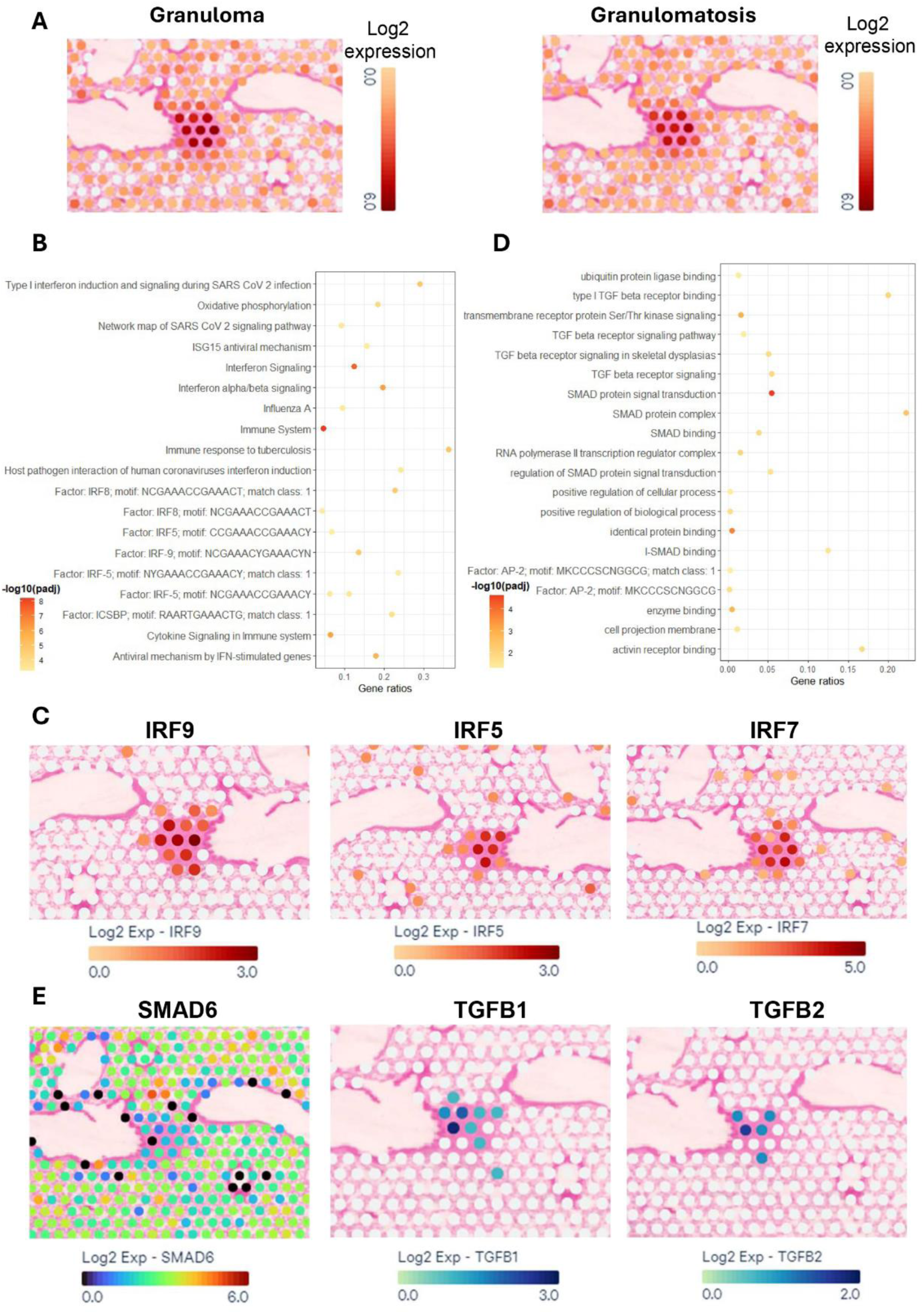
Gene expression changes and pathway analysis of lymphoid aggregate. **A)** Aggregated gene expression of granuloma and granulomatosis gene sets from MSigDB projected on hematoxylin stain of lungs from coinfection mice. Spot-wise gene expression aggregate was projected on lymphoid cell aggregate and its surround tissue. Color key represents aggregate of expression values of all the genes in the granuloma gene set (DOCK2, CYBC1, PIK3CG, CYBB, IRF8, NF1, DNASE2, and ACP5) and granulomatosis gene set (DOCK2, CYBC1, PIK3CG, CYBB, IRF8, NF1, DNASE2, and ACP5). **B)** Top 20 pathways based on an adjusted p value identified in over representation analysis on gene upregulated in cluster 13 spanning lymphoid cell aggregate in coinfection mice. Gene ratios were computed as the percentage of total DEGs in the given pathway term. **C)** Expression levels of interferon regulatory factors enriched among upregulated genes in lymphoid cell aggregate were projected individually on lymphoid cell aggregate and its surround tissue. **D)** Top 20 pathways identified in over representation analysis on gene down regulated in cluster 13 as shown like figure 4B. **E)** Down regulation of SMAD6, an inhibitory SMAD and upregulation of TGFB1/2 in lymphoid cell aggregate. Gene expression levels quantified in spatial transcriptomics analysis were projected on hematoxylin-stained tissue section. A, C, E) Color key represents aggregate/individual gene expression values in log2 scale.

Encouraged by these results, to elucidate the molecular changes occurring in the early stages of granuloma development, we performed pathway analysis to identify signaling pathways enriched in the lymphoid cell aggregate region. We performed differential expression analysis using the Seurat ‘find-markers’ function to identify genes that were up- or downregulated in Cluster 13. Applying a log2FC cutoff of 2 and an adjusted p value of 0.05, we identified 383 upregulated and 20 downregulated genes. Pathway analysis was performed independently for both the up- and downregulated genes, revealing significant enrichment of pathways related to immune response to viral, *Mtb*, and bacterial infections, as well as B-cell and NK cell activity, cytokine signaling, glucose metabolism, oxidative phosphorylation, ROS generation, interferon signaling, and the activation of several interferon regulatory factor (IRF) transcription factors in genes upregulated in the lymphoid cell aggregate (**Figure 6B, Supplementary Table 1**). Interestingly, oxidative phosphorylation and ROS generation are crucial for pathogen clearance within granulomas^31^. Transcriptional targets of several interferon regulatory factors (IRF5, IRF7, and IRF9) were enriched in upregulated genes (**Figure 6B-C, Supplementary Table 1**). Additionally, type I interferon signaling promotes *Mtb* infection by facilitating the release of neutrophil extracellular traps^32^ and suppressing interferon gamma and interleukin 1α/β signaling^33^.

Intriguingly, similar analysis of downregulated genes revealed enrichment of multiple pathways related to downregulation of inhibitory SMAD expression in lymphoid cell aggregates, suggesting potential activation of TGFβ signaling (**Figure 6D**, **Supplementary Table 2**). This finding was corroborated by decreased expression of SMAD6, which inhibits TGFβ signaling, alongside upregulation of TGFβ signaling ligands, namely, TGFβ1/2, in the same tissue region, suggesting activated TGFβ signaling (**Figure 6E**). Taken together, these results suggest that lymphoid cell aggregates present several granuloma-like characteristics prior to evolving into mature granulomas.

### IRF7 and TGFβ signaling are active in T cells and macrophages

To determine the immune cell types that harbor active IRF7 and TGFβ signaling in the lungs, we performed immunofluorescence staining on lung sections from mice infected with *Mtb* alone or coinfected with *Mtb* and HIV. We used cell-specific markers along with IRF7 and TGFβ antibodies to assess whether these signaling molecules colocalize with immune cells. Given that granulomas are predominantly composed of macrophages, and HIV targets T cells, we examined these cell types for active IRF7 and TGFβ signaling. Interestingly, expression levels of macrophage marker CD68 were decreased in *Mtb*/HIV-coinfected mice compared to those infected with *Mtb* alone, suggesting that HIV infection may impair macrophage recruitment and survival in the lungs, consistent with findings by Lepard et al., 2022^14^ (Figure 7A). Similarly, the signal levels of T cells marker CD3 were comparatively lower in the coinfected group relative to the *Mtb*-only group, likely reflecting the progressive depletion of CD4⁺ T cells driven by HIV infection (Figure 7B), consistent with findings from Berges et al., 2006^34^. In addition, we observed consistent colocalization of intracellular markers IRF7 and TGFβ with these cells in both *Mtb*-only and *Mtb*/HIV-coinfected lungs (Figure.7A&B). These results indicate that HIV coinfection reduces the overall infiltration of macrophages and T cells in the lungs while these cells maintain active IRF7 and TGFβ signaling.

**Figure 7.**
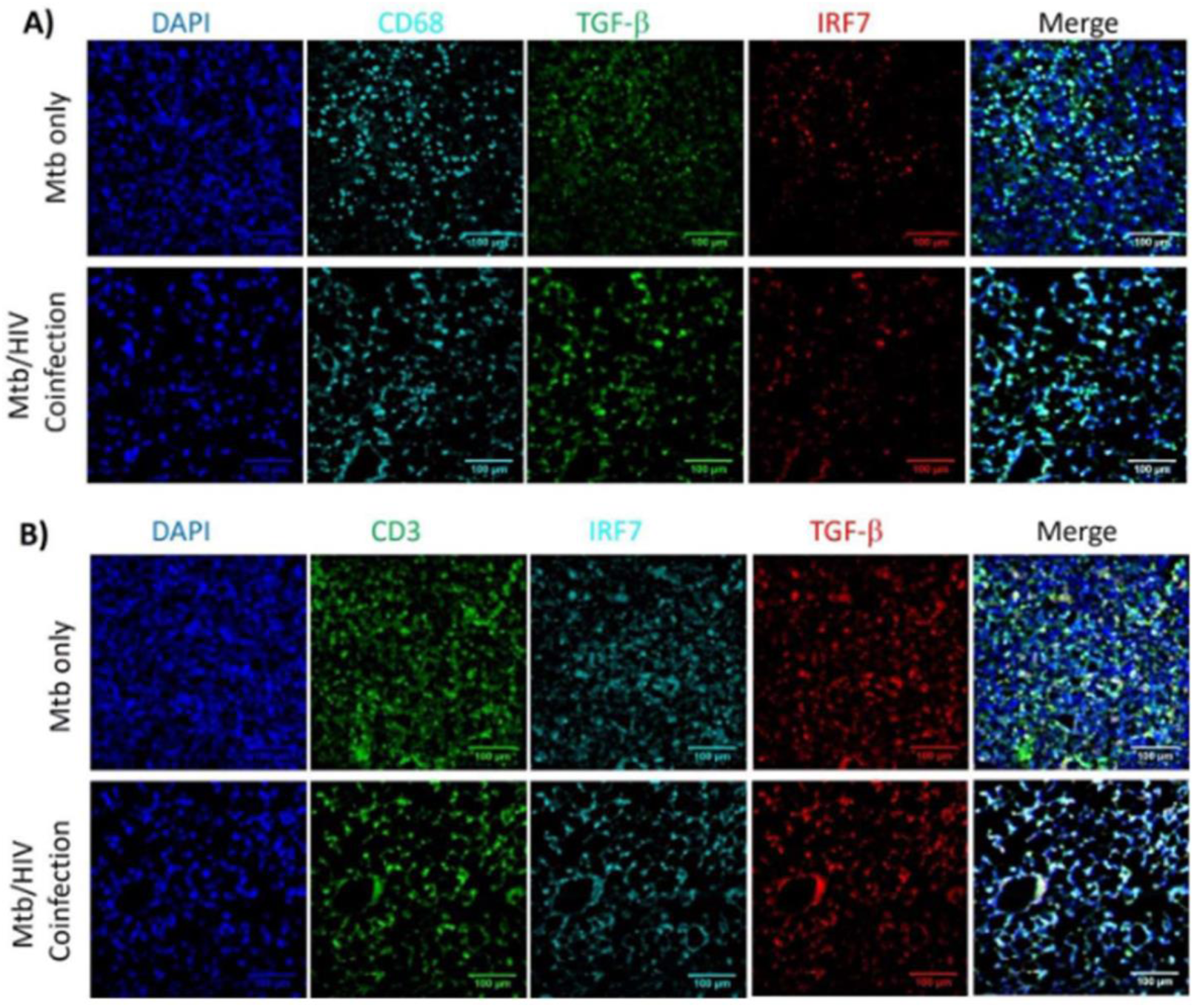
**Dynamics of macrophages and T cells in the lungs of mice infected with *Mtb* or *Mtb*/HIV coinfection**. Immunofluorescence staining was performed on FFPE lung sections from mice infected with *Mtb* alone or coinfected with *Mtb* and HIV to examine immune cell infiltration and the colocalization of intracellular signaling molecules IRF-7 and TGF-β. **A)** Sections were surface-stained for human macrophage marker CD68, along with intracellular markers IRF-7 and TGF-β. **B)** Sections were surface-stained for human T-cell marker CD3, along with intracellular IRF-7 and TGF-β. DAPI-supplemented mounting medium was used for nuclear staining. Representative images were acquired at 20× magnification using a Zeiss LSM-500 confocal microscope. Scale bars: 100 μm.

## Discussion

Novel spatial transcriptomic technologies provide a powerful tool to assess molecular and immunological changes in situ in various tissues. This approach provides deeper insights into the host response to infection, and the disease development or resistance^35,36^. In this study, we employed these technologies to evaluate early molecular changes induced by HIV, *Mtb*, and their coinfection in humanized mouse lung tissues. Although the lung is the main target organ of *Mtb*, it is also profoundly impacted by HIV-induced generalized immunosuppression^9^ due to massive depletion of the CD4^+^ T cell population^8,9^. Following *Mtb* single infection, increased CD4+ T cell migration to affected sites aids in pathogen control^4^. Moreover, severe local depletion of this CD4+ T cell subset in HIV-infected hu-mice (with or without *Mtb* coinfection) is a hallmark of HIV that has been previously reported^8^ and observed in the present work. The approach used in the present study also revealed alterations in less-studied populations, notably nonclassical monocytes and NK cells. Intriguingly, while HIV and *Mtb* single infection increased NK cell populations compared to uninfected controls, coinfection had the opposite effect. This pattern aligns with reported changes in certain phenotypes of NK cells in HIV/*Mtb*-coinfected patients, suggesting that alterations in this cell subset may vary in accordance with the phenotypes present in the individual subject^37^.

A considerable portion of the differences evidenced in the cell populations presented within the tissue corresponded not only to infection but also to distinct histological regions, with bronchi and major airways clearly distinguished from interstitial tissue. However, even within clustered airway regions, viral and bacterial infections altered the phenotypical landscape. This is evidenced by the high presence of alveolar macrophages expressing CCL3 in the airway regions from all the infected animals, in contrast with the uninfected controls. CCL3 aids in alveolar macrophage activation and efforts to control bacterial and viral pathogens^26,38^. Recent studies have also shown increased CCL3 levels in COVID-19 patients with alveolar damage, underscoring the important role of these cells in pulmonary immunity^39^.

The reduction in M1 macrophages, neutrophils, and classical monocytes in HIV and coinfected airways likely creates a permissive environment for *Mtb* survival. This supports the hypothesis that HIV coinfection disrupts immune cell infiltration patterns necessary for controlling *Mtb*, thereby exacerbating TB pathogenesis. Conversely, the increased abundance of NK cells and innate lymphoid cells type 2 (ILC2) in HIV and coinfected airways suggests a shift toward innate lymphoid-driven immunity. This shift may reflect an attempt to compensate for the loss of adaptive immunity (e.g., CD4⁺ T cell depletion) or contribute to immunopathology, such as excessive tissue remodeling driven by ILC2.

The finding of a structurally organized and clearly differentiated cell cluster with a unique transcriptomic profile in a sample from an *Mtb*/HIV-coinfected mouse provides insight into the early phenotypical and signaling changes induced by this dual infection. Spatial analysis revealed the presence of both lymphoid and myeloid cells in the lymphoid aggregates. Remarkably, despite HIV-induced generalized CD4^+^ T cell depletion in the tissue, CD4+ T cells was found within the aggregate, primarily in the inner region, alongside B cells and plasmacytoid DCs. Notably, unlike mature granulomas, which typically feature a necrotic core surrounded by macrophages and epithelioid cells with lymphocytes in the outer layer^2,3,7^, the lymphoid aggregate in the *Mtb*/HIV-coinfected tissue lacked necrosis, with monocytic cells positioned in the outer layer. This is likely because this cell aggregate is in the early stages of forming a granulomatous structure, and the time after infection has not yet been sufficient to generate necrosis. This observation is consistent with previous findings in a murine *Mtb* model, where inflammatory cell aggregates, mainly adjacent to the circulation or bronchi, was detected as early as 2 weeks post-infection, with the first necrotic changes detected approximately one month after infection^27,40^. Moreover, our results show that the transcriptomic alterations found in this nascent granuloma-like structure mirror those reported for mature granulomas, reinforcing the notion that the lymphoid aggregate found in this sample is in the early stages of granuloma development.

The process of granuloma establishment involves precise modulation of the interferon pathway, mainly through the upregulation of multiple IRFs, as shown in the cells present in this granuloma-like structure. Some of the IRF genes that are upregulated in the lymphoid aggregate in this study have been previously reported to be crucial for granuloma formation and maintenance, as evidenced by IRF8-deficient mice, which fail to form these structures and control *Mtb* growth after infection^41^. However, the increased transcription of different IRF genes suggests that the IFN response is regulated via multiple mechanisms, including cell differentiation and innate immunity. This likely stems from the initial induction of type-I IFNs and activation of the IFN pathway and IFN-stimulated genes (ISGs) after infection. However, the relevance of this multifaceted IFN modulation in more advanced stages of structural remodeling remains to be elucidated. While some studies suggest that mature granulomas (after 9 weeks post-infection) do not show upregulation of these pathways, others report increased IRF transcription in these structures, particularly in nonimmune cell types (fibroblasts)^6,42^.

Consequently, the activation of the immune response by the IFN pathway leads to increased energy requirements in the immune cells present in the lymphoid aggregate, as exemplified by the upregulation of oxidative phosphorylation-related genes. This metabolic shift has been a well-documented consequence of HIV, *Mtb*, and *Mtb*/HIV coinfection^43–45^. Our results show that the generalized metabolic changes induced by *Mtb*/HIV coinfection also occur locally in the affected tissues and, particularly in the affected areas, starting at an early stage of infection.

Our findings indicate that, unlike the regulation of the IFN pathway, TGFβ regulation is absent. TGFβ inhibition of *Mtb*-specific CD4^+^ T cells contributes to the maintenance of *Mtb* within mature granuloma structures in rhesus macaque and mouse models^46^. Additionally, TGFβ and its associated pathways have also been proposed as determinants of myeloid cell differentiation in TB granulomas^47^, whereas computational modeling suggests enhanced bacterial clearance in the absence of TGFβ^48^. The concurrent reduction in macrophages and T cells, coupled with active IRF7 and TGFβ signaling, suggests a complex interplay where HIV coinfection may skew and disrupt immune responses. Investigating whether IRF7 drives pro-inflammatory responses or TGFβ promotes immunosuppression could provide insights into disease progression.

### Conclusions

The modulation of the immune response by pathogens is an important determinant of the outcome of infection, ranging from generalized immunosuppression, as seen with HIV, to more nuanced modulation of the immune response induced by *Mtb*. This study examines the effects of these pathogens, whether in the context of single infection or coinfection, on lung tissue in a humanized NSG-SGM3-IL15 mouse model at an early timepoint after inoculation. Our findings suggest that early infection triggers alterations in cellular infiltration and structural remodeling of the tissue, mirroring the patterns observed in humans and other animal models. These findings further validate the humanized NSG-SGM3-IL15 mouse strain as a reliable animal model for studying HIV, *Mtb*, and *Mtb*/HIV coinfection. Nevertheless, future studies using higher-resolution spatial transcriptomics will further elucidate the transcriptomic changes in lung immune cell populations at the single-cell level, providing further detail into specific changes in different cell populations, both in terms of phenotype and signaling. Such insights may pave the way for novel intervention strategies that improve infection outcomes in patients infected with HIV and *Mtb* infections.

## Data sharing statement

All data supporting the findings of this study are available in the manuscript. Raw and processed spatial transcriptome data was deposited in the National Center for Biotechnology Information (NCBI) Gene Expression Omnibus (GEO)and can be accessed at GSE303811. If there are any special requests or questions for the data, please contact the corresponding author (G.Y.).

## Acknowledgments

We thank the technical support from the Cancer Prevention and Research Institute of Texas (CPRIT RP240610). We thank the Texas Advanced Computing Center (TACC) at The University of Texas at Austin for providing computational resources that have contributed to the research results reported within this paper. We’d like to thank Drs. Buka Samten and Amy Tvinnereim for their aid in *Mtb* infection of the humanized mice.

## Contributions

Sitaramaraju Adduri: Conceptualized the study, designed and performed the experiments, analyzed data, edited figures, and wrote the manuscript.

Jose Alejandro Bohorquez: Conceptualized the study, designed and perform the experiments, analyzed data, edited figures, and wrote the manuscript.

Rajesh Mani: Perform the experiments, analyzed data, edited figures, and wrote the manuscript. Omoyeni Adejare: Perform the experiments.

Diego Rincon: Analyzed data.

Joshua K. Kleam: Perform the experiments.

Mounika Duggineni: Perform the experiments.

Andy Omeje: Perform the experiments.

Torry Tucker: Edited the manuscript, provided critical resources.

Nagarjun V Konduru: Designed the experiments, edited the manuscript.

Guohua Yi: Conceptualized and guided the study, secured the funding, designed the experiments, analyzed data, edited figures, and wrote, edited, and finalized the manuscript.

