## Supplementary Materials for "Spatial transcriptomic analysis of HIV and tuberculosis coinfection in a humanized mouse model reveals unique transcription patterns, immune responses and early morphological alterations"

### Supplementary Figures:

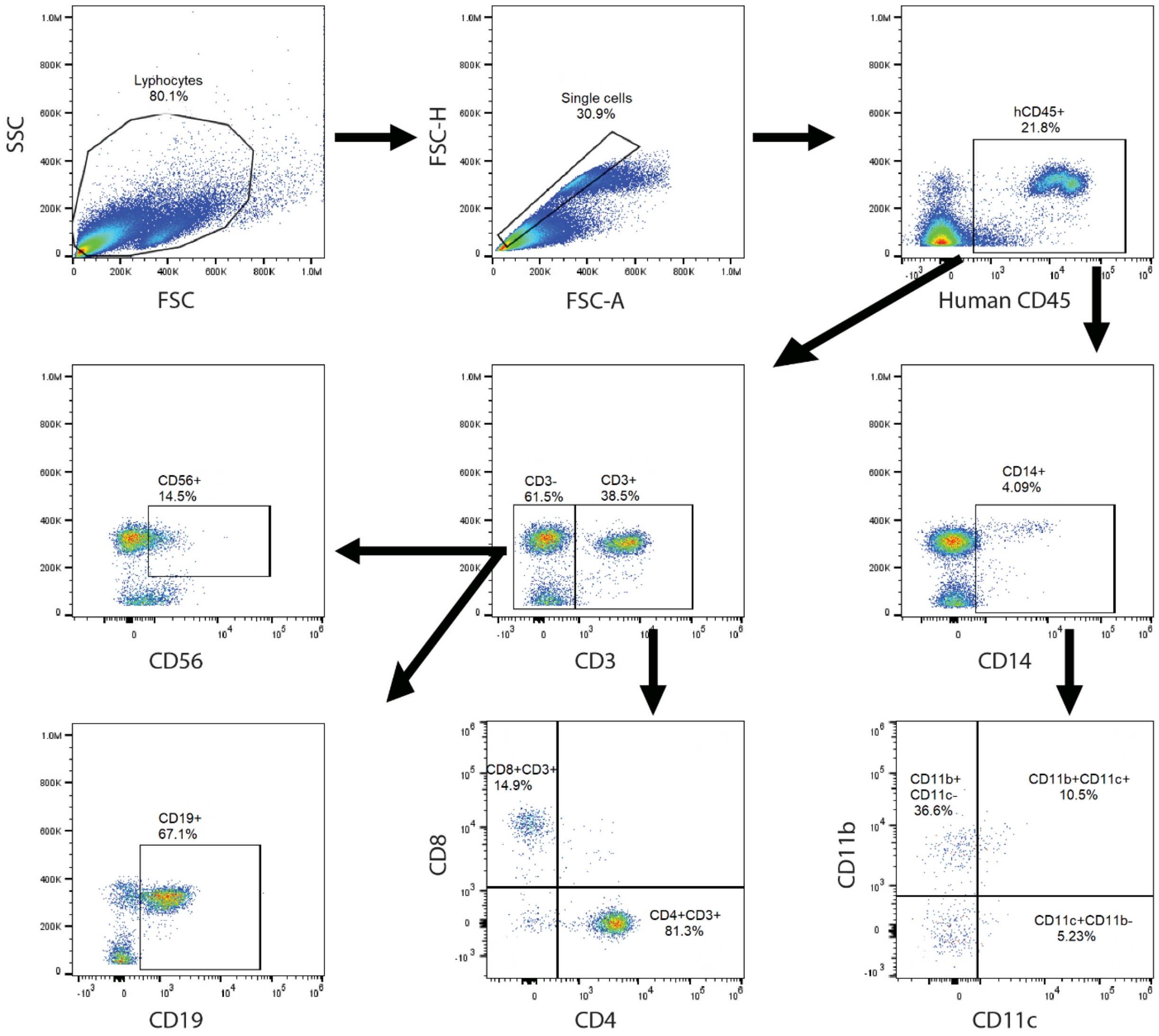


**Figure S1: Engraftment and differentiation of human immune cells in humanized NSG-SGM3-IL15 mice.** A dose of 2 x $10^5$ human CD34+ cord blood-derived hematopoietic stem cells was injected into each 100 cGy-irradiated 4-5-week-old NSG-SGM3-IL15 mice. After three months of humanization, the stem cell engraftment and the human immune cell differentiation were characterized staining the PBMCs with fluorescent anti-human antibodies specific for human CD45 (human cells), CD3 (T cells), CD4 (CD4+ T cells), CD8 (CD8+ T cells), CD14 (monocytes), CD11b (macrophages), CD11c (Dendritic cells), CD19 (B cells), and CD56 (NK cells), and the stained cells were counted by flow cytometry.

**Figure S2**

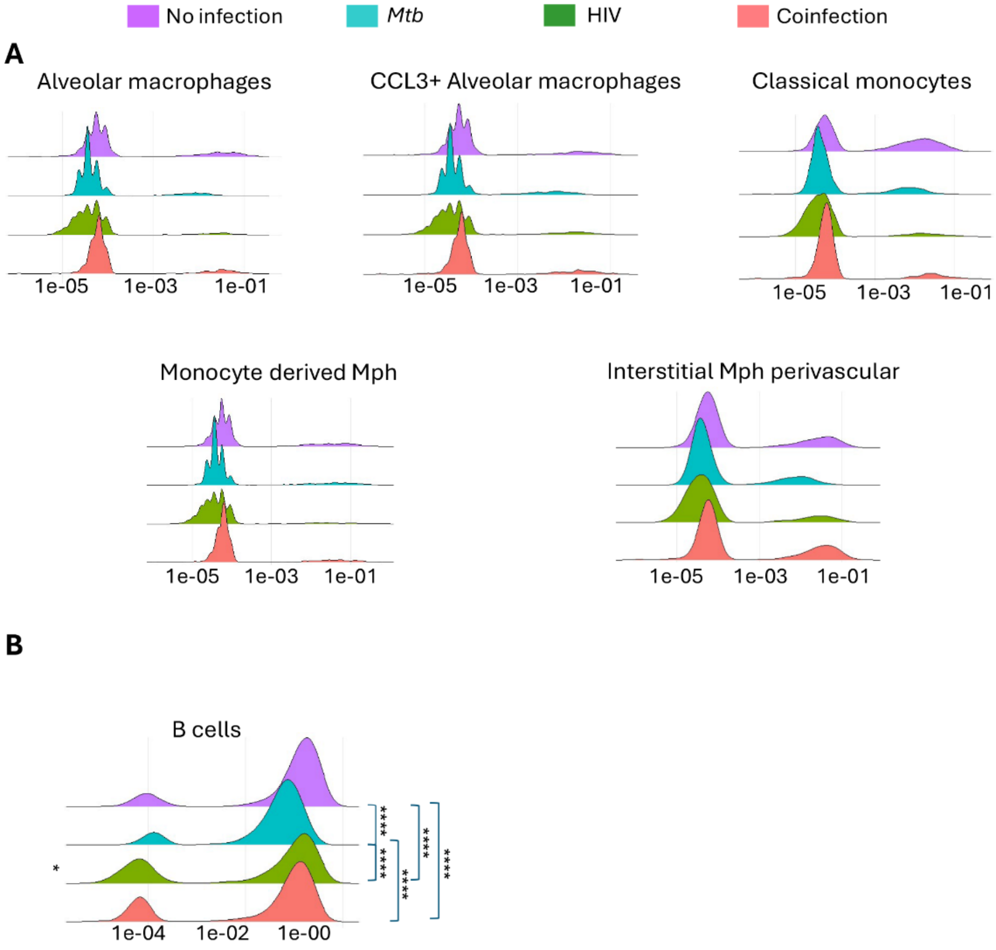


**Figure S2:** Cell type weights for each spot in the samples based on RCTD analysis were plotted for each infection type. X axis shows cell type weights which are proportional to the cell type abundance in the tissue.

Figure 3

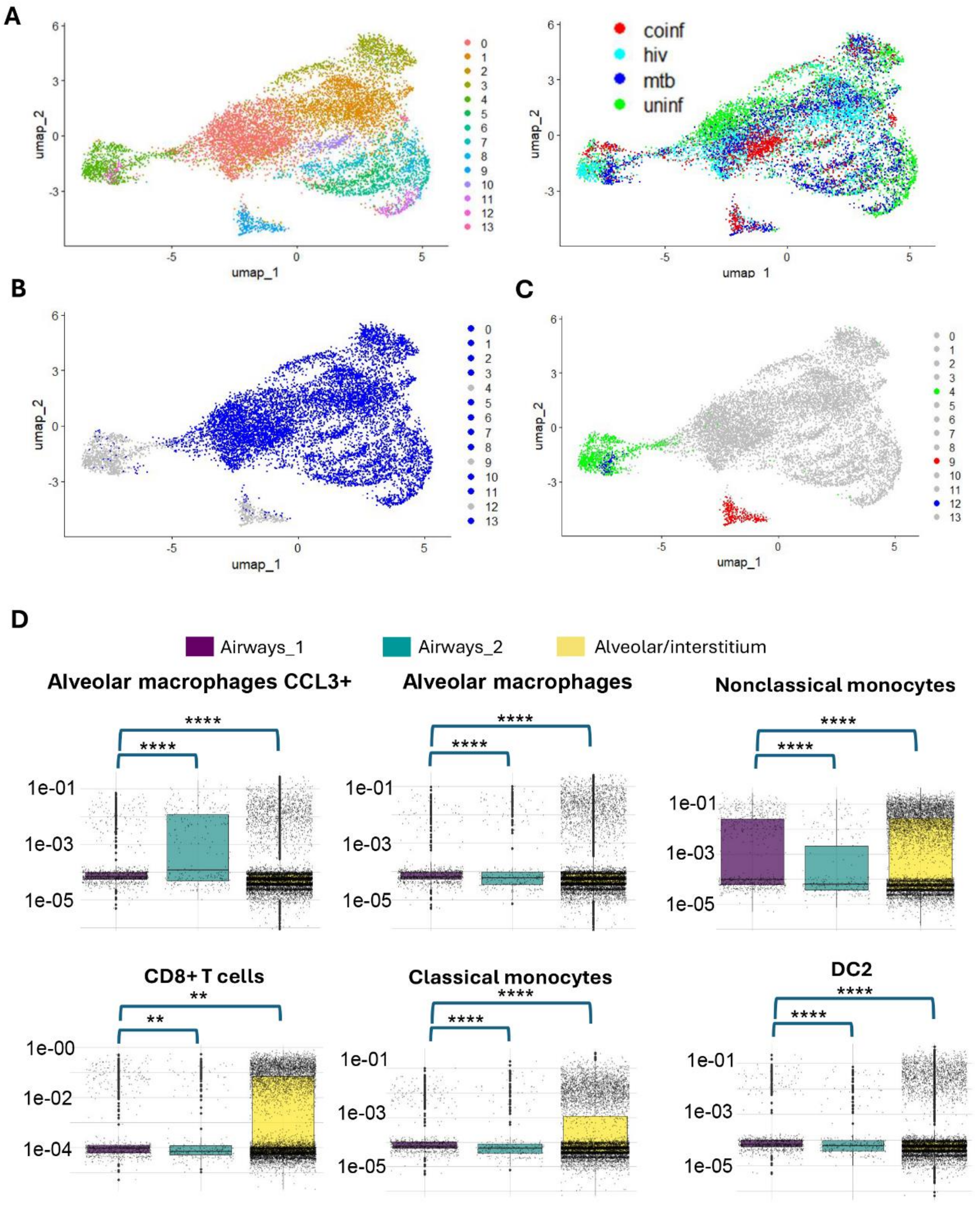


**Figure S3:** Spatial distribution map of expression of CD4 in lungs from *Mtb* infection and coinfection mice. Ir-Intercalator was used to stain nucleus. 195Pt_ICSK2 and 160Gd_CD66b were used for staining plasma membrane and granulocytes, respectively.

Figure S4

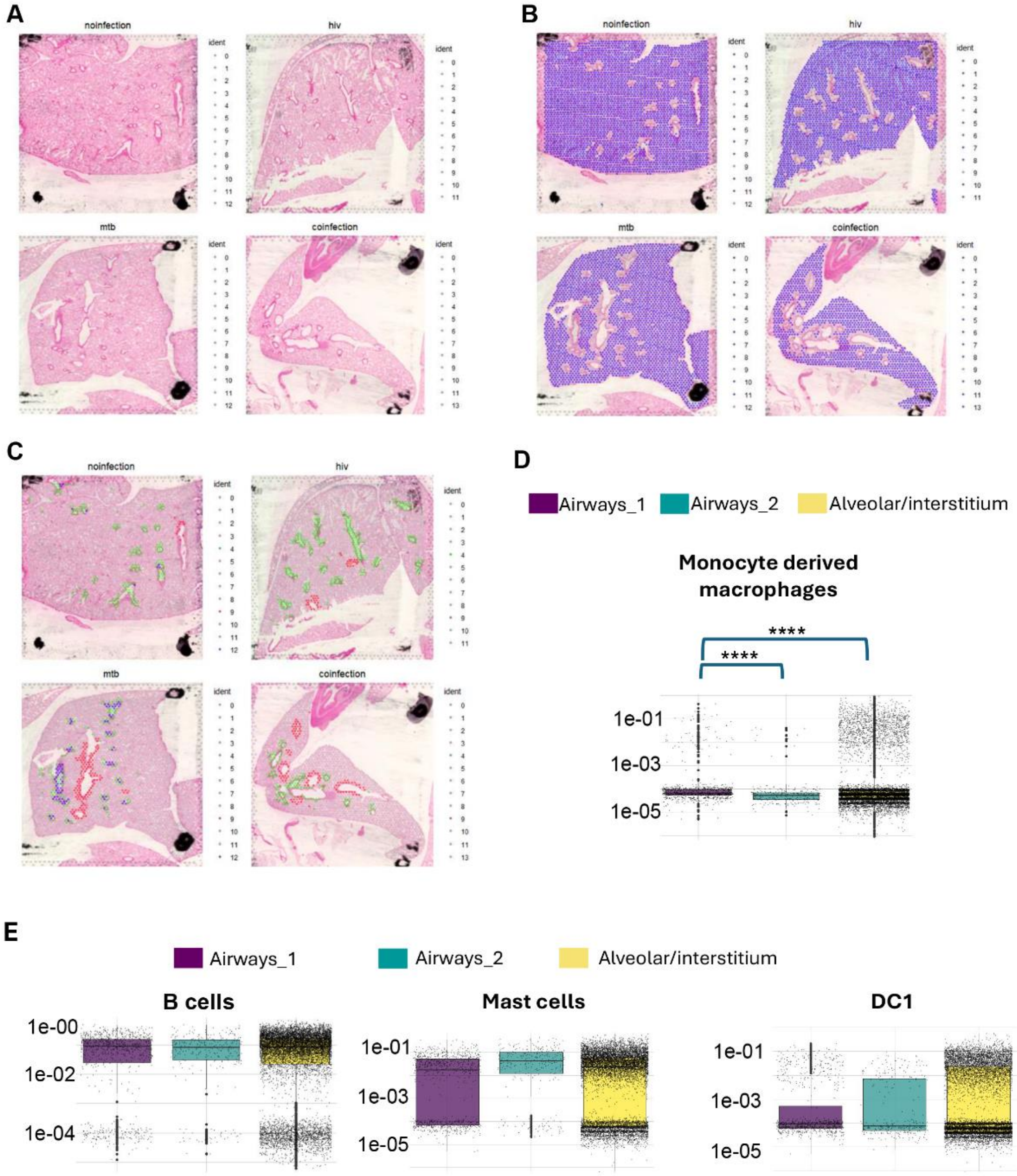


**Figure S4:** Clusters identified based on gene expression in all four samples are shown along with hematoxylin staining of corresponding tissue. 12 clusters were detected in each of infection, *Mtb*, and coinfection lungs. 11 clusters were detected in HIV lungs. Cluster 13 was detected exclusively in coinfection lungs. B-C) Spots corresponding to alveolar/interstitium cluster (shown in blue) (B), airway_1 (shown in blue/green) (C) and airway-2 (shown in red) (C) clusters projected on hematoxylin staining of corresponding tissue. D-E) Cell type weights obtained in RCTD deconvolution analysis were plotted for each cluster. Y axis shows cell type weights which are proportional to the cell type abundance in the corresponding cluster. The color key indicated the cluster identifier. The difference in cell weight distributions was computed using Kruskal Wallis test followed by Dunn's test for pairwise comparisons. P values were adjusted for multiple hypothesis testing using Bonferroni correction method. Significance levels are indicated as *$P$ < .05, **$P$ < .01, ***$P$ < .001, and ****$P$ < .0001.

Figure S5

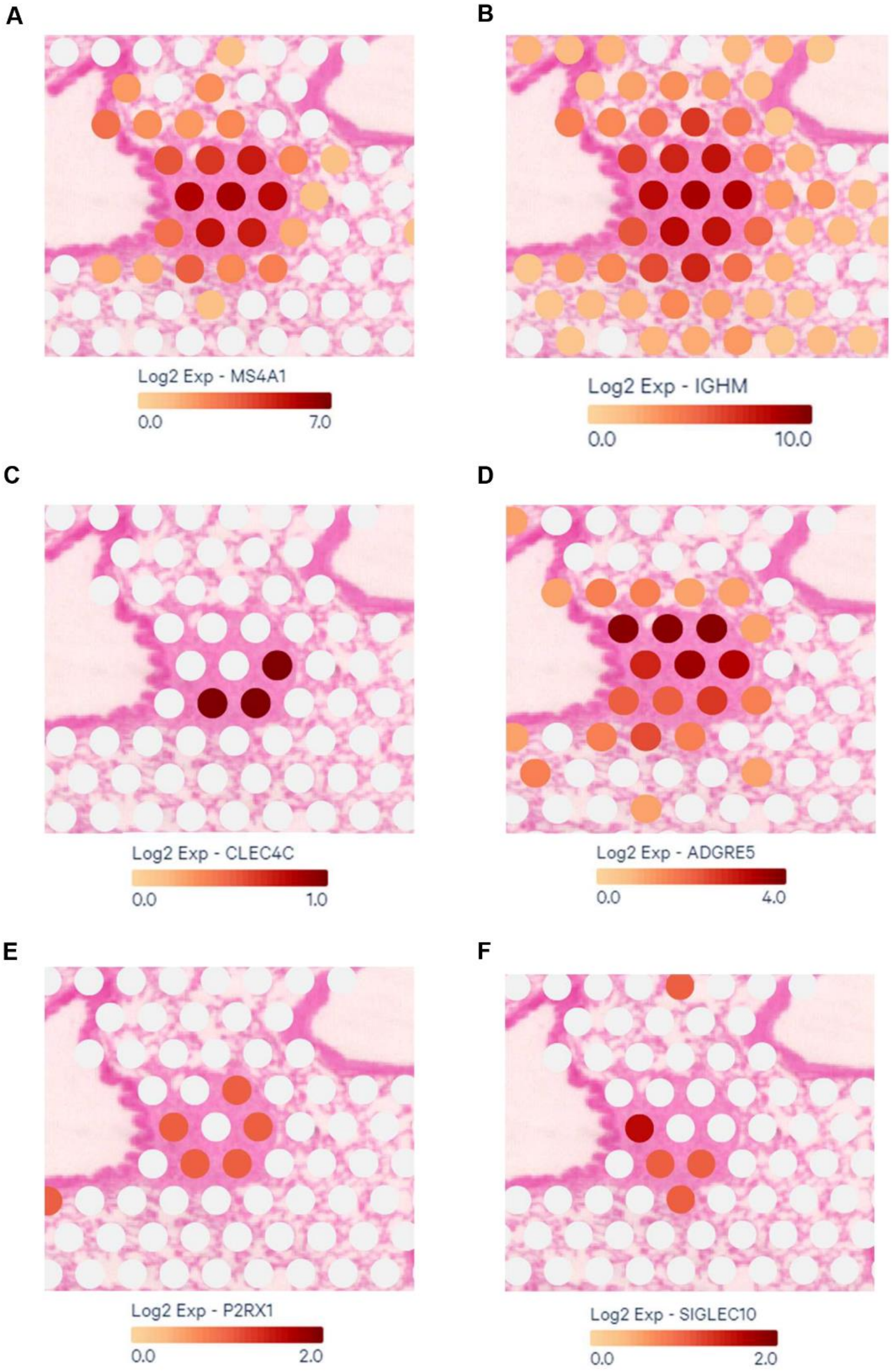


**Figure S5:** Spatial map of expression of A) CD20 (MS4A1), B) IgM (IGHM), C) CD303 (CLEC4C), D) CD97 (ADGRE5), E) P2RX1, and F) Siglec10 within and around the lymphoid aggregate. Spot-wise gene expression data was projected on hematoxylin staining image of the lung tissue. Color key represents the level of transcript expression in log2 scale.

### Figure S6

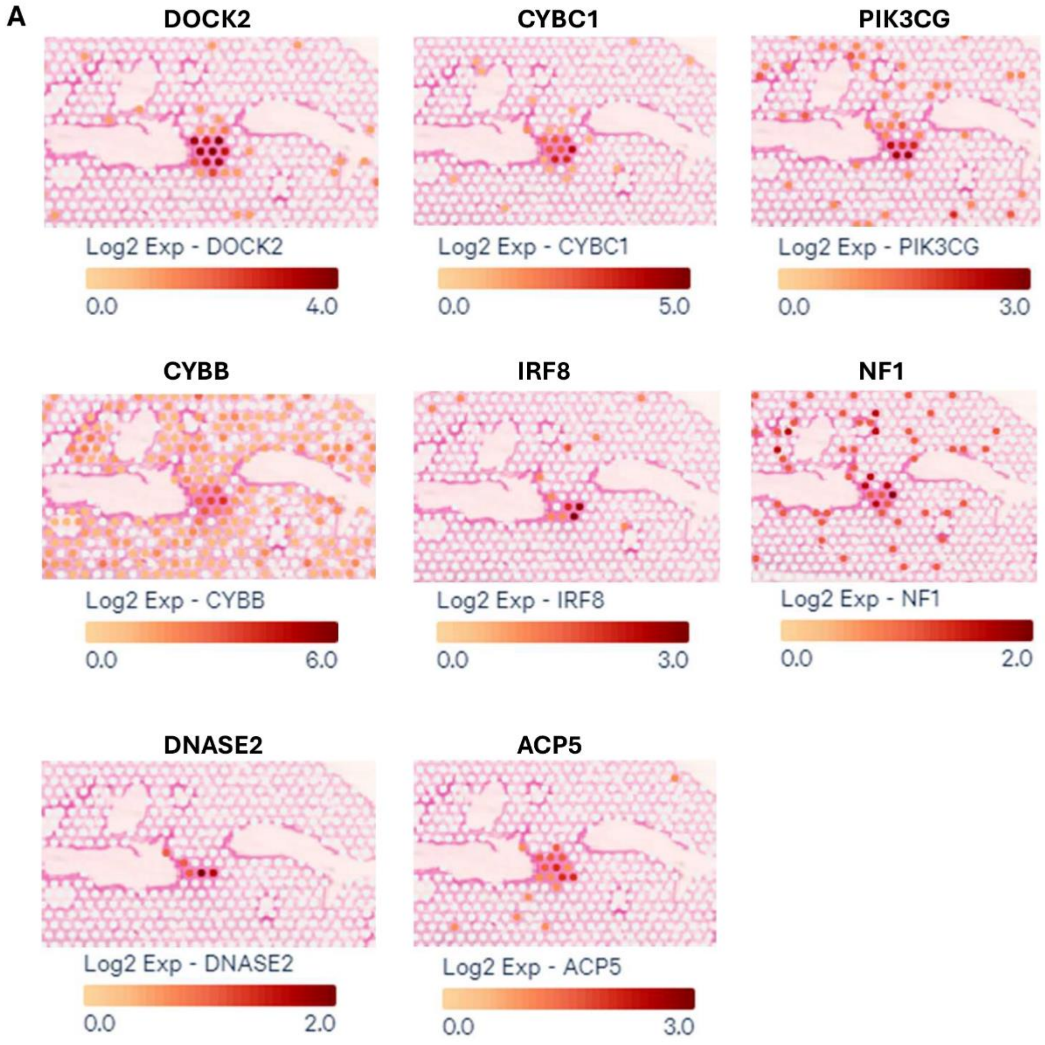


**Figure S6: Expression pattern of genes belonging to granuloma gene set in lung tissue of coinfection mice quantified in spatial transcriptomic analysis.** A) Expression levels of genes in HP_GRANULOMA were plotted individually. Spot-wise gene expression was projected on lymphoid cell aggregate and its surround tissue. Color key represents aggregate of expression values in log2 scale.

### Figure S7

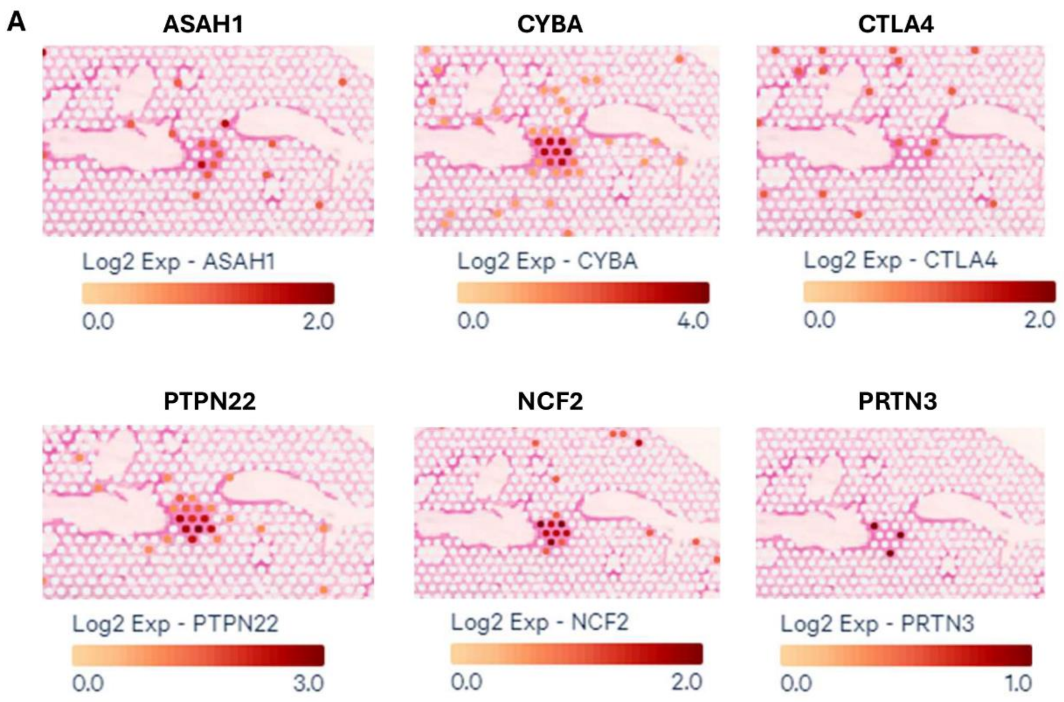


**Figure S7: Expression pattern of genes belonging to granulomatosis gene set in lung tissue of coinfection mice quantified in spatial transcriptomic analysis.** A) Expression level of genes in HP_ GRANULOMATOSIS gene set were plotted individually. Spot-wise gene expression was projected on lymphoid cell aggregate and its surround tissue. Color key represents aggregate of expression values in log2 scale. Spatial transcriptome analysis did not quantify the expression of the following genes belonging to Granulomatosis gene set: HLA-DPA1, HLA-DPB1, and NCF1. Expression pattern of CYBB, a member of granulomatosis gene was shown in Figure S6.

**Supplementary Tables:**

**Table 1. Pathway terms enriched among the upregulated genes spots spanning lymphoid aggregate detected in coinfection lung.**

| Term Name | Term Id | Negative Log10 of Adjusted P Value | Term Size | Query Size | Intersection Size |
|---|---|---|---|---|---|
| **Immune System** | REAC:R-HSA-168256 | 8.210 | 2052 | 268 | 97 |
| **Interferon Signaling** | REAC:R-HSA-913531 | 7.544 | 193 | 268 | 24 |
| **Interferon alpha/beta signaling** | REAC:R-HSA-909733 | 6.137 | 71 | 268 | 14 |
| **Cytokine Signaling in Immune system** | REAC:R-HSA-1280215 | 6.063 | 719 | 268 | 46 |
| **Antiviral mechanism by IFN-stimulated genes** | REAC:R-HSA-1169410 | 5.575 | 78 | 268 | 14 |
| **Immune response to tuberculosis** | WP:WP4197 | 4.935 | 22 | 224 | 8 |
| **Type I interferon induction and signaling during SARS CoV 2 infection** | WP:WP4868 | 4.802 | 31 | 224 | 9 |
| **Factor: IRF8; motif: NCGAAACCGAAACT; match class: 1** | TF:M04020_1 | 4.716 | 44 | 370 | 10 |
| **Factor: IRF-9; motif: NCGAAACYGAAACYN** | TF:M11681 | 4.664 | 103 | 370 | 14 |
| **Oxidative phosphorylation** | WP:WP623 | 4.026 | 60 | 224 | 11 |
| **Factor: ICSBP; motif: RAARTGAAACTG; match class: 1** | TF:M00699_1 | 3.805 | 41 | 370 | 9 |
| **Factor: IRF-5; motif: NCGAAACCGAAACY** | TF:M11672 | 3.529 | 126 | 370 | 14 |
| **Network map of SARS CoV 2 signaling pathway** | WP:WP5115 | 3.525 | 217 | 224 | 20 |
| **Factor: IRF-5; motif: NCGAAACCGAAACY** | TF:M11671 | 3.420 | 398 | 370 | 25 |
| **Influenza A** | KEGG:05164 | 3.412 | 167 | 211 | 16 |
| **Host pathogen interaction of human coronaviruses interferon induction** | WP:WP4880 | 3.409 | 33 | 224 | 8 |
| **ISG15 antiviral mechanism** | REAC:R-HSA-1169408 | 3.318 | 70 | 268 | 11 |
| **Factor: IRF8; motif: NCGAAACCGAAACT** | TF:M04020 | 3.308 | 1019 | 370 | 44 |

| Term Name | Term Id | Negative Log10 of Adjusted P Value | Term Size | Query Size | Intersection Size |
|---|---|---|---|---|---|
| **Factor: IRF-5; motif: NYGAAACCGAAACY; match class: 1** | TF:M11670_1 | 3.303 | 34 | 370 | 8 |
| **Factor: IRF5; motif: CCGAAACCGAAACY** | TF:M04016 | 3.277 | 322 | 370 | 22 |
| **Cell Cycle** | REAC:R-HSA-1640170 | 3.130 | 677 | 268 | 38 |
| **Factor: IRF-9; motif: NYGAAACYGAAACYN** | TF:M11680 | 3.112 | 136 | 370 | 14 |
| **The citric acid (TCA) cycle and respiratory electron transport** | REAC:R-HSA-1428517 | 3.084 | 177 | 268 | 17 |
| **Cell Cycle, Mitotic** | REAC:R-HSA-69278 | 3.084 | 547 | 268 | 33 |
| **Parkinson disease** | KEGG:05012 | 3.024 | 265 | 211 | 20 |
| **Factor: IRF-7; motif: NCGAAANCGAAANYN** | TF:M11679 | 3.002 | 634 | 370 | 32 |
| **Respiratory electron transport, ATP synthesis by chemiosmotic coupling, and heat production by uncoupling proteins.** | REAC:R-HSA-163200 | 2.865 | 127 | 268 | 14 |
| **Factor: IRF-8; motif: NCGAAACYGAAACYN; match class: 1** | TF:M11685_1 | 2.813 | 123 | 370 | 13 |
| **Factor: E2F-1; motif: TTTSGCGCGMNR; match class: 1** | TF:M00516_1 | 2.760 | 2126 | 370 | 71 |
| **Oxidative phosphorylation** | KEGG:00190 | 2.548 | 134 | 211 | 13 |
| **Factor: ISGF-3; motif: CAGTTTCWCTTTYCC** | TF:M00258 | 2.501 | 839 | 370 | 37 |
| **Epstein-Barr virus infection** | KEGG:05169 | 2.473 | 198 | 211 | 16 |
| **Hepatitis C** | KEGG:05160 | 2.413 | 158 | 211 | 14 |
| **Measles** | KEGG:05162 | 2.412 | 138 | 211 | 13 |
| **Factor: IRF-5; motif: NYGAAACCGAAACY** | TF:M11670 | 2.394 | 883 | 370 | 38 |
| **Diabetic cardiomyopathy** | KEGG:05415 | 2.366 | 202 | 211 | 16 |
| **Electron transport chain OXPHOS system in mitochondria** | WP:WP111 | 2.359 | 104 | 224 | 12 |
| **Prion disease** | KEGG:05020 | 2.350 | 271 | 211 | 19 |
| **Resolution of Sister Chromatid Cohesion** | REAC:R-HSA-2500257 | 2.307 | 123 | 268 | 13 |
| **Retinoblastoma gene in cancer** | WP:WP2446 | 2.297 | 89 | 224 | 11 |

| Term Name | Term Id | Negative Log10 of Adjusted P Value | Term Size | Query Size | Intersection Size |
|---|---|---|---|---|---|
| **Thermogenesis** | KEGG:04714 | 2.179 | 232 | 211 | 17 |
| **Factor: ICSBP; motif: RAARTGAAACTG** | TF:M00699 | 2.025 | 1213 | 370 | 46 |
| **Factor: Rb:E2F-1:DP-1; motif: TTTSGCGC; match class: 1** | TF:M00740_1 | 2.015 | 4732 | 370 | 126 |
| **Type II interferon signaling** | WP:WP619 | 2.004 | 37 | 224 | 7 |
| **Amyotrophic lateral sclerosis** | KEGG:05014 | 1.999 | 363 | 211 | 22 |
| **Bacterial invasion of epithelial cells** | KEGG:05100 | 1.919 | 77 | 211 | 9 |
| **Chemical carcinogenesis - reactive oxygen species** | KEGG:05208 | 1.899 | 221 | 211 | 16 |
| **RHO GTPases Activate Formins** | REAC:R-HSA-5663220 | 1.835 | 136 | 268 | 13 |
| **Cell cycle** | KEGG:04110 | 1.832 | 157 | 211 | 13 |
| **Measles virus infection** | WP:WP4630 | 1.828 | 136 | 224 | 13 |
| **Mitotic Prometaphase** | REAC:R-HSA-68877 | 1.799 | 199 | 268 | 16 |
| **B cell receptor signaling pathway** | KEGG:04662 | 1.747 | 81 | 211 | 9 |
| **Interferon type I signaling pathways** | WP:WP585 | 1.747 | 54 | 224 | 8 |
| **Hepatitis B** | KEGG:05161 | 1.696 | 162 | 211 | 13 |
| **Factor: IRF7; motif: NCGAAARYGAAANT** | TF:M04018 | 1.680 | 628 | 370 | 29 |
| **Alzheimer disease** | KEGG:05010 | 1.679 | 382 | 211 | 22 |
| **Huntington disease** | KEGG:05016 | 1.667 | 305 | 211 | 19 |
| **Respiratory electron transport** | REAC:R-HSA-611105 | 1.666 | 103 | 268 | 11 |
| **Interferon gamma signaling** | REAC:R-HSA-877300 | 1.595 | 87 | 268 | 10 |
| **Natural killer cell mediated cytotoxicity** | KEGG:04650 | 1.578 | 124 | 211 | 11 |
| **Non genomic actions of 1 25 dihydroxyvitamin D3** | WP:WP4341 | 1.548 | 73 | 224 | 9 |
| **Extrafollicular and follicular B cell activation by SARS CoV 2** | WP:WP5218 | 1.502 | 74 | 224 | 9 |
| **Factor: IRF-8; motif: NCGAAACYGAAACYN** | TF:M11685 | 1.501 | 1672 | 370 | 56 |
| **Factor: IRF-2; motif: NAANYGAAASYR; match class: 1** | TF:M08775_1 | 1.492 | 2790 | 370 | 82 |
| **Factor: STAT2; motif: RRGRAANNGAAACTGAAAN** | TF:M10080 | 1.491 | 352 | 370 | 20 |

| Term Name | Term Id | Negative Log10 of Adjusted P Value | Term Size | Query Size | Intersection Size |
|---|---|---|---|---|---|
| **Complex I biogenesis** | REAC:R-HSA-6799198 | 1.465 | 57 | 268 | 8 |
| **Mitotic Spindle Checkpoint** | REAC:R-HSA-69618 | 1.439 | 109 | 268 | 11 |
| **Amplification of signal from unattached kinetochores via a MAD2 inhibitory signal** | REAC:R-HSA-141444 | 1.386 | 92 | 268 | 10 |
| **Amplification of signal from the kinetochores** | REAC:R-HSA-141424 | 1.386 | 92 | 268 | 10 |
| **Factor: Ncx; motif: NAATNAATTAATAANWW; match class: 1** | TF:M01420_1 | 1.344 | 2157 | 370 | 67 |
| **Cell Cycle Checkpoints** | REAC:R-HSA-69620 | 1.324 | 289 | 268 | 19 |
| **Factor: ATF-2; motif: TTACGTAA; match class: 1** | TF:M00040_1 | 1.316 | 6522 | 370 | 159 |
| **Factor: IRF; motif: NNGAAANTGAAANN; match class: 1** | TF:M08887_1 | 1.304 | 79 | 370 | 9 |
| **Factor: E2F; motif: TTTSGCGSG** | TF:M00939 | 1.303 | 6627 | 370 | 161 |

**Table 2. Pathway terms enriched among the downregulated genes spots spanning lymphoid aggregate detected in coinfection lung.**

| Term Name | Term Id | Adjusted p value | Negative Log10 of Adjusted P Value | Term Size | Query Size | Intersection Size |
|---|---|---|---|---|---|---|
| **SMAD protein signal transduction** | GO:0060395 | 2.27E-05 | 4.644 | 92 | 19 | 5 |
| **Identical protein binding** | GO:0042802 | 1.81E-04 | 3.743 | 2162 | 19 | 11 |
| **Transmembrane receptor protein serine/threonine kinase signaling pathway** | GO:0007178 | 9.68E-04 | 3.014 | 374 | 19 | 6 |
| **Enzyme binding** | GO:0019899 | 1.47E-03 | 2.833 | 2103 | 19 | 10 |
| **Heteromeric SMAD protein complex** | GO:0071144 | 3.05E-03 | 2.516 | 8 | 19 | 2 |
| **SMAD protein complex** | GO:0071141 | 3.91E-03 | 2.407 | 9 | 19 | 2 |
| **RNA polymerase II transcription regulator complex** | GO:0090575 | 9.55E-03 | 2.020 | 258 | 19 | 4 |
| **Type I transforming growth factor beta receptor binding** | GO:0034713 | 9.92E-03 | 2.003 | 10 | 19 | 2 |
| **TGF beta receptor signaling** | WP:WP560 | 1.20E-02 | 1.922 | 55 | 14 | 3 |
| **SMAD binding** | GO:0046332 | 1.36E-02 | 1.868 | 78 | 19 | 3 |
| **activin receptor binding** | GO:0070697 | 1.45E-02 | 1.838 | 12 | 19 | 2 |
| **TGF beta receptor signaling in skeletal dysplasias** | WP:WP4816 | 1.48E-02 | 1.831 | 59 | 14 | 3 |
| **Factor: AP-2; motif: MKCCCSCNGGCG** | TF:M00189 | 1.76E-02 | 1.754 | 10911 | 19 | 19 |
| **Positive regulation of biological process** | GO:0048518 | 1.86E-02 | 1.731 | 6250 | 19 | 15 |
| **Regulation of SMAD protein signal transduction** | GO:0060390 | 2.53E-02 | 1.596 | 57 | 19 | 3 |
| **I-SMAD binding** | GO:0070411 | 2.64E-02 | 1.579 | 16 | 19 | 2 |
| **Cell projection membrane** | GO:0031253 | 3.30E-02 | 1.481 | 356 | 19 | 4 |
| **Term Name** | **Term Id** | **Adjusted p value** | **Negative Log10 of Adjusted P Value** | **Term Size** | **Query Size** | **Intersection Size** |

| | | | | | | |
|---|---|---|---|---|---|---|
| **Transforming growth factor beta receptor signaling pathway** | GO:0007179 | 4.41E-02 | 1.355 | 206 | 19 | 4 |
| **Ubiquitin protein ligase binding** | GO:0031625 | 4.53E-02 | 1.344 | 308 | 19 | 4 |
| **Positive regulation of cellular process** | GO:0048522 | 4.61E-02 | 1.336 | 5717 | 19 | 14 |
| **Factor: AP-2; motif: MKCCCSCNGGCG; match class: 1** | TF:M00189_1 | 4.82E-02 | 1.317 | 6273 | 19 | 15 |
